# Predicting the prevalence of complex genetic diseases from individual genotype profiles using capsule networks

**DOI:** 10.1101/2022.12.13.520248

**Authors:** Xiao Luo, Xiongbin Kang, Alexander Schönhuth

**Affiliations:** Life Science & Health, Centrum Wiskunde & Informatica, Amsterdam, The Netherlands; Genome Data Science, Faculty of Technology, Bielefeld University, Bielefeld, Germany

## Abstract

Diseases that have a complex genetic architecture tend to suffer from considerable amounts of genetic variants that, although playing a role in the disease, have not yet been revealed as such. Two major causes for this phenomenon are genetic variants that do not stack up effects, but interact in complex ways; in addition, as recently suggested, the omnigenic model postulates that variants interact in a holistic manner to establish disease phenotypes.

We present DiseaseCapsule, as a capsule network based approach that explicitly addresses to capture the hierarchical structure of the underlying genome data, and has the potential to fully capture the non-linear relationships between variants and disease. DiseaseCapsule is the first such approach to operate in a whole-genome manner when predicting disease occurrence from individual genotype profiles.

In experiments, we evaluated DiseaseCapsule on amyotrophic lateral sclerosis (ALS) and Parkinson’s disease (PD), with a particular emphasis on ALS because known known to have a complex genetic architecture, so being affected by considerable missing heritability (40%). On ALS, DiseaseCapsule achieves 86.9% accuracy on held out test data in predicting disease occurrence, thereby outperforming all other approaches by large margins. Also, DiseaseCapsule required sufficiently less training data for reaching optimal performance. Last but not leaset, the systematic exploitation of the network architecture yielded 922 genes of particular interest, and 644 ”non-additive” genes that are crucial factors in DiseaseCapsule, but have no effect within linear schemes.

## Introduction

Amyotrophic lateral sclerosis (ALS) is a rare primary neurodegenerative syndrome characterized by human motor system degeneration. The incidence of ALS in the European population is about 2.16 per 100,000 person-years [1]. As the disease progresses, predominant symptoms are amyosthenia, paralysis and ultimately death. After the onset of symptoms, 50% of the patients die within 2.5 years, with further 20% of the patients dying within 5 to 10 years [2]. Recent studies show that NAD^+^ replenishment can improve clinical features of ALS patients, indicating an encouraging potential novel treatment for ALS [3, 4]. To date, ALS is still not curable, but symptomatic treatment can significantly improve life quality and survival of the affected [5]. Because symptoms tend to be insidious and escape their identification, diagnosis of ALS often comes at a considerable delay: it is common that it requires 12 months for a diagnosis to be definitive [6]. Therefore, most patients miss the advantageous opportunities of early intervention [7].

The elusiveness of the disease during its early stages and the corresponding delays in establishing diagnosis explain why efficient methods and tools for predicting prevalence and occurrence of ALS have life-saving potential. Employing such tools when symptoms are still inconclusive supports practitioners in taking measures sufficiently early. In addition to their value in clinical practice, predictive models can help to elucidate the pathological mechanisms of ALS. This, eventually, supports the development of optimally targeted therapies.

Various studies, for example on the heritability of sporadic ALS in twin pairs [8], the SOD1 mutation in mouse models [9], or the largest GWAS of ALS to date [10], have demonstrated that ALS is a complex disorder that has an encompassing genetic background [11]. At the same time, next-generation sequencing technology has meant a considerable boost for genome-wide association studies (GWAS). The central purpose of GWAS is to reveal the correlation of (in particular disease) phenotypes with individual genetic make-ups. However, the genetic variants delivered by GWA type studies have been amounting to only 10% of the heritability [12] of ALS. This contrasts non GWA type (e.g. twin record based) studies that have evaluated the heritability of ALS to amount to 50% [13]. It is therefore reasonable to assume that the missing heritability of ALS amounts to approximately 40%.

The striking amount of missing heritability, or even the idea that missing heritability is an ill-posed concept as soon as genotype-phenotype-relationships are supposed to be based on the omnigenic [14, 15], and not the polygenic model are supposed to be the major reasons for the poor prognosis of ALS. In summary, the major methodological challenges are: *first*, the association signals of complex diseases can spread across most of the genome instead of involving just a few core pathways [14, 16], that is the omnigenic models applies. Effects of associated variants are often weak, so do not exceed the stringent significance thresholds that apply in GWAS [16, 17]. *Second*, the linear models that underlie GWAS analysis techniques cannot detect non-additive genetic effects. Epistasis, where mutations condition each other in non-additive, functional relationships [17, 18] cannot be uncovered by standard GWAS techniques, for example. Several studies have modeled gene-gene interactions [19, 20, 21]. However, only a few of them have been directly applied to predicting the prevalence of complex genetic diseases.

Methods that aim to exploit non-additive relationships have left behind various open questions. When being based on statistical hypothesis testing, they tend to suffer from a lack of power: Because the number of hypotheses tested exceed the ones of traditional approaches by orders of magnitude, the necessary correction for multiple testing masks significant signals. On the other hand, approaches that are based on—potentially omnigenic—machine learning models are relatively rare, and so far have left ample room for improvements.

The fact that polygenic risk scores, as a purely linear approach to predict occurrence or prevalence of diseases, tend to underperform is further evidence for the insufficiency of additive schemes. Last but not least, most recent work presented how to assess the association of ALS with entire linkage disequilibrium (LD) blocks, and not just single variants [22]. There, LD blocks were modeled in terms of a Bayesian hidden variable based framework that combines single variants within the blocks in a non-linear manner. For prioritizing LD blocks, epigenetic information was used. Consequently, this study revealed various new variants associated with ALS that linear approaches had been blind to.

Deep learning, as a predominant machine learning approach, has established the state-of-the-art in areas such as image classification, face detection, speech recognition, and many more. The basis of deep learning are feedforward neural networks that stack many hidden layers of neurons on top of each other; the depth of such a network is defined to be the number of such hidden layers.

Extensions of the universal approximation theorem [23] support the stacking of layers in particular [24], where the accuracy of the prediction increases exponentially on increasing numbers of hidden layers. These extensions provide a theoretical basis for the insight that not only the width of the layers, but the depth of the network is crucial for reaching superior levels of accuracy; from a historical perspective, raising the depth of neural networks lead to the breakthrough advantages they are enjoying today [25]. The intuitive idea that motivates the shape and arrangement of such networks is to detect and arrange patterns in a hierarchical way: subpatterns, detected by the early layers, form overarching patterns detected by later layers. Overall, this leads to elevated levels of resolution when mapping the data [26].

Convolutional neural networks (CNN’s) reflect network architectures that are particularly suitable to implement the idea of hierarchies of patterns [27]. While CNN’s such as AlexNet [25], VGG Net [28], ResNet [29] and DenseNet [30], indeed achieved the most striking breakthrough successes in deep learning, major criticisms with regard to interpretability (“deep black boxes”) [31] and the enormous demand for training data for reaching superior performance [32] had been remaining. Both these points are highly relevant in clinical applications [33, 34]. While collecting clinical data can be expensive and demanding, the inability to explain can raise ethical concerns.

Capsule networks (CapsNets) [35, 36] were presented as a remedy for addressing such critical issues. The major motivation for CapsNets was to classify distorted or entangled patterns in images correctly. The improved modeling of spatial hierarchies, as empowered by the “viewpoint invariance property”, led to major improvements with respect to the accuracy of the predictions, as the driving issue. Beyond that, the ”viewpoint invariance property” is the likely reason for the reduced requirements in terms of training data that were observed in comparison with CNN’s: thanks to this property, capsule networks can generalize to recognizing objects regardless of the angle relative to which the corresponding training data was presented, which implies that less training data is required. Further, although not primarily intended, CapsNets also enabled a human-mind friendly interpretation of results. Here, following these ideas, we will demonstrate that the architecture of CapsNets can be used to not only associate genes with the disease in general, but also to highlight relevant genes interacting in non-additive ways in particular.

Therefore, CapsNets have shown to have the potential to resolve two issues of primary concern in biomedical applications. Recent studies indicate that the potential of CapsNets to learn complex hierarchical structures can indeed be leveraged also for biomedical data. Successful examples are biological networks (see e.g. [37] for a review), or single cell classification [38], among further applications. These applications of CapsNets add to the earlier (innumerous) applications of ordinary CNN’s or machine learning models in biology or medicine, for example for cancer prognosis and prediction [39], for obesity classification [40], or for mechanistic exploration and drug development in neurodegenerative diseases [41, 42, 43]. Despite the above mentioned issues for standard CNN’s remaining, they point out the great potential of deep learning in biomedical applications in general.

As particularly related work, there also have been recent applications of deep learning when predicting phenotype from genotype. Relevant examples are the detection of epistasis [44, 45], or the prediction of ALS from single nucleotide polymorphism (SNP) array based genotype profiles of the promoter regions from 4 particularly selected chromosomes [46]. Last but not least, the non-linear approach mentioned earlier [22] models non-linearities within the range of LD blocks. However, subsequently, the effects of LD blocks are summarized in a linear scheme, which fails to model non-linear interactions across LD blocks. In a brief summary of the current status of related work, we find approaches that model non-linear interactions across promoter regions in 4 selected chromosomes [46], on the one hand, or within LD blocks, on the other hand, where, however, LD blocks are confined to interact linearly across the genome.

The omnigenic model [14] requires to model gene-gene interactions across the entire genome, in a way that allows genes to interact in non-additive ways for establishing their joint effects. From this point of view, none of the approaches presented so far in the literature builds on the omnigenic model as its foundation. Either, non-linear interactions can only be captured in the range of LD blocks [22], or, when captured across the range of LD blocks, only promoter regions or selected chromosomes are considered [46].

Our approach will be the first approach to model whole-genome wide, non-additive interactions between genes. For that, it takes a route that one can consider the opposite of the protocol presented in [22]: we summarize local (gene-range) effects of variants linearly, and then combine the local effects in non-additive ways globally (across the entire genome). Arguably, we present the first *deep* neural network based approach that caters to the omnigenic model as a conceptual basis. More specifically, from a methodical point of view, we present the first approach that employs capsule networks, as an advanced deep neural network class of functions, to map genotypes onto phenotypes. In that regard, we recall that the universal approximation theorem and its extensions [23, 24] support that neural networks can capture arbitrary, and in particular non-additive, functional relationships.

## Results

We present *DiseaseCapsule*, as a framework that can handle genome-scale variant input and reveal non-additive interactions across genes. In the following, first, we briefly summarize our approach, which consists of two basic stages: while the first stage refers to reducing the dimensionality of the (enormous, because genome-wide) input according to a new data reduction protocol, the second stage picks up the condensed data, and runs it through a capsule network that has been designed to predict phenotypes of interest from individual genotype profiles.

Although DiseaseCapsule establishes a model that is generic with respect to the genetic disease it can predict, we focus on the occurrence / prevalence of ALS as phenotype in the following. The reasons for this are the availability of the excellent, sufficiently large amounts of data that were raised in the frame of Project MinE (https://www.projectmine.com). Most importantly, we emphasize ALS, based on the afore-mentioned large missing heritability. For analogous reasons, also the earlier related studies [22, 46] rely on data raised in the frame of Project MinE. See Methods for exact descriptions of the data.

In a brief summary of the results that follow (see Table 1), DiseaseCapsule outperforms all prior approaches when predicting occurrence/prevalence of ALS from individual whole-genome genotype profiles by large margins. DiseaseCapsule achieves 86.9% accuracy, establishing a relative increase of 40% over prior standards, such as polygenic risk scores or various other possible state-of-the-art approaches. In this, DiseaseCapsule seems to be particularly superior in terms of recall: evidently, capsule networks are able to recognize the genetic patterns that drive the disease in a more sensitive way.

**Table 1.** Classification results for ALS test data. The values are represented as percentages. MLP: multilayer perceptron, SVM: support vector machine, CNN: convolutional neural network. *a,b,c,d* represent PRS-based models that the SNPs were selected by GWAS with the threshold (*P* < 5 × 10^-2^, *P* < 5 × 10^-4^, *P* < 5 × 10^-6^ and *P* < 5 × 10^-8^) respectively. *Note*: The best score is marked in bold.

| Dimension reduction | Classifier | Accuracy | Precision | Recall | F1-Score |
| --- | --- | --- | --- | --- | --- |
| Gene-PCA | DiseaseCapsule | **86.9** | 85.2 | 89.4 | **87.2** |
| Gene-PCA | MLP | 84.2 | 92.2 | 74.8 | 82.6 |
| Gene-PCA | Logistic Regression | 78.2 | 71.1 | 94.8 | 81.3 |
| Gene-PCA | SVM | 76.3 | **94.8** | 55.8 | 70.3 |
| Gene-PCA | CNN | 74.5 | 86.1 | 58.5 | 69.7 |
| Gene-PCA | Random Forest | 63.3 | 73.0 | 42.1 | 53.4 |
| Gene-PCA | AdaBoost | 62.7 | 86.3 | 30.2 | 44.7 |
| All-PCA | DiseaseCapsule | 81.9 | 80.7 | 83.8 | 82.2 |
| All-PCA | Logistic Regression | 78.1 | 70.7 | **96.0** | 81.4 |
| All-PCA | SVM | 76.3 | **94.8** | 55.8 | 70.3 |
| All-PCA | MLP | 72.5 | 85.2 | 54.4 | 66.4 |
| All-PCA | AdaBoost | 67.6 | 84.8 | 42.9 | 57.0 |
| All-PCA | Random Forest | 64.1 | 73.3 | 44.4 | 55.3 |
| All-PCA | CNN | 53.8 | 54.8 | 42.5 | 47.9 |
| - | Polygenic Risk Score<sup>a</sup> | 81.8 | 91.5 | 70.2 | 79.4 |
| - | Polygenic Risk Score<sup>b</sup> | 78.5 | 84.4 | 69.8 | 76.4 |
| - | Polygenic Risk Score<sup>c</sup> | 74.2 | 76.8 | 69.4 | 72.9 |
| - | Polygenic Risk Score<sup>d</sup> | 63.5 | 63.5 | 63.3 | 63.4 |

Results will further demonstrate how to reveal 922 core genes that are crucial for the capsule network based classification process. Interestingly, the majority of these 922 genes play documented roles in the nervous system, but many of which have not been associated with ALS so far (only 13 genes can be found in the ALSoD database). Investigating this further, we provide a list of 644 genes that play a crucial role during capsule network based classification (so interacting in complex ways), while not raising significant signals in linear schemes. While it still remains to be explored why these “non-additive” genes are crucial for identifying the occurrence (or non-occurrence) of ALS, it is clear already that standard GWAS type approaches will remain blind to them, without raising cohorts of patients to unrealistic sizes.

### Workflow

DiseaseCapsule implements a two-step approach to successfully deal with genome-scale variant screens. The first step consists of a novel protocol to perform dimensionality reduction. This protocol enables to capture millions of polymorphic loci in a way that enables subsequent application of non-additive methods. The second step then is the application of capsule networks, as a fundamentally non-additive model, to the reduced data. These two steps are preceded by basic quality control and batch effect removal, as well as a basic step to filter out variants that do not matter (regardless of the underlying approach). See Fig. 1 for an illustration. In the following, we describe the essence of the two steps, and refer the reader to the Methods section for full methodical details. In the following, the data on which DiseaseCapsule and competing methods are validated refers to 10,405 DNA array based, whole-genome genotype samples from the Dutch cohort of Project MinE.

**Fig. 1.**
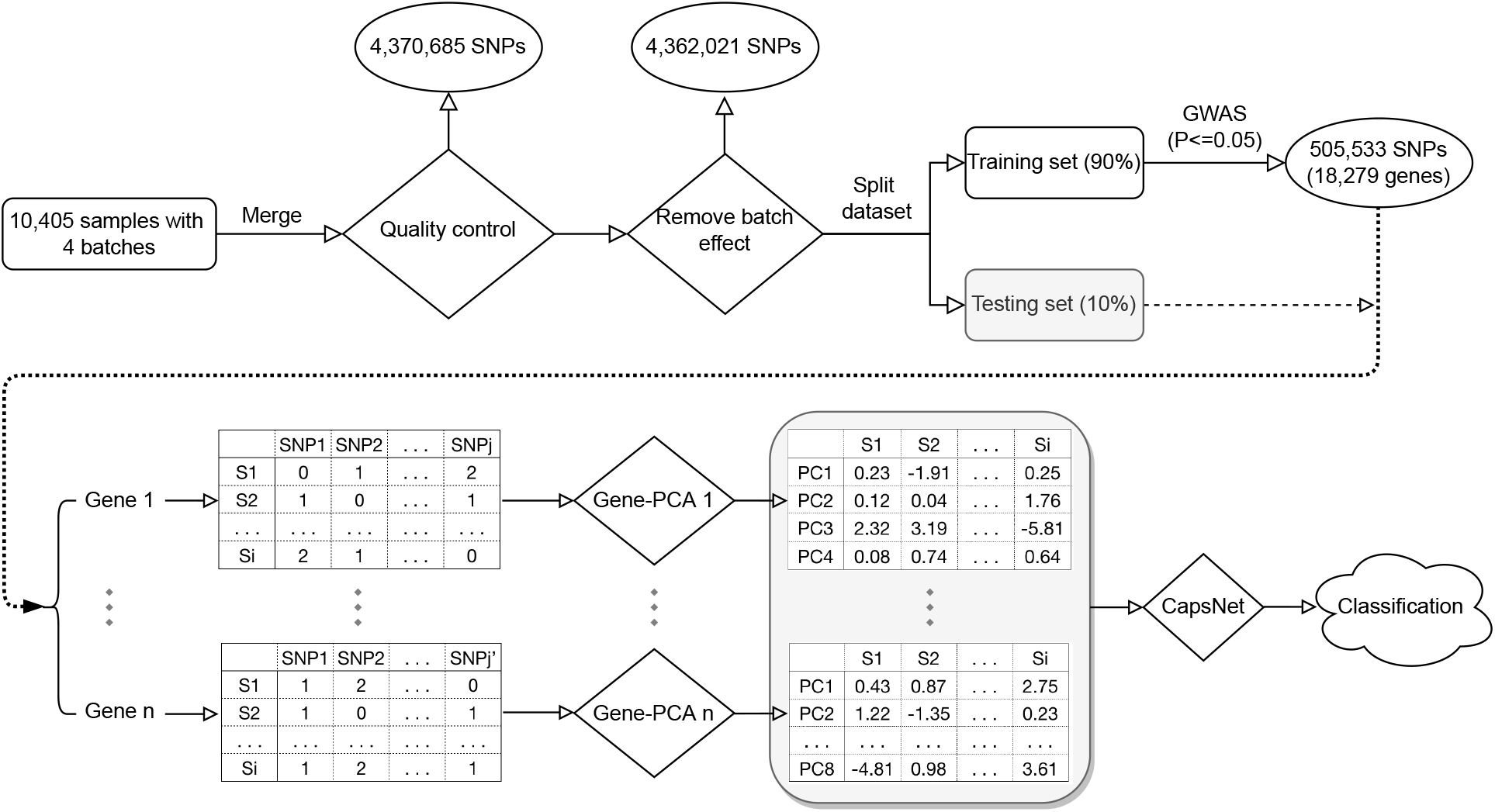
An overview of the workflow. In the tables for each gene, *S*_1_, *S*_2_,…, *S_i_* represent sample IDs, and *PC*_1_, *PC*_2_ ,…, *PC_k_* represent the 1, 2,…, *k*-th principal component of each Gene-PCA, respectively. The number of *PC* is 8 or 4 or 1, which depends on the length of the input, i.e. the number of SNPs.

### Dimensionality Reduction

#### Challenge

Global, genome-wide application of principal component analysis (PCA) consistently reduces the dimensionality of the input data. When both PCA and association models are additive in nature, they effectively combine.

The use of non-linear / non-additive association models, however, becomes annihilated by prior application of *genome-wide* PCA. Because principal components reflect linear combinations of variants, subsequent application of non-linear techniques yields non-additive combinations of whole genome wide, additive combinations of variants. In principle, this still has the potential to model certain genome-wide interactions non-linearly. However, transiting from variants themselves to genome-wide linear combinations of them, and only then applying non-additive techniques does not only confuse the situation conceptually, but also annihilates the advantages of the non-linear approach in practice. The latter is documented by our results (see Tables 1 and 3 “All-PCA” for related results).

**Table 2.** Test accuracy, precision and recall using non-additive genes for prediction. Models are retrained only using 644 non-additive genes.

| Model | Accuracy | Precision | Recall |
| --- | --- | --- | --- |
| Gene-PCA+DiseaseCapsule | 0.712 | 0.715 | 0.706 |
| Gene-PCA+Logistic Regression | 0.512 | 0.522 | 0.292 |
| Difference | 0.200 | 0.193 | 0.414 |

**Table 3.** Testing accuracy, precision, recall and F1 score for different cohorts. C5 & C44 and C1 & C3 were used for trainning and testing respectively. The values are represented as percentages. *This model is PRS-based that the SNPs were selected by GWAS with the threshold (*P* < 5 × 10^-2^). *Note*: The best score is marked in bold.

| Dimension reduction | Classifier | Accuracy | Precision | Recall | F1-Score |
| --- | --- | --- | --- | --- | --- |
| Gene-PCA | DiseaseCapsule | **81.6** | 85.8 | 71.3 | **77.9** |
| Gene-PCA | MLP | 74.5 | 94.3 | 46.5 | 62.3 |
| Gene-PCA | LogisticRegression | 72.8 | 64.5 | 88.7 | 74.7 |
| Gene-PCA | CNN | 65.3 | 82.7 | 29.6 | 43.6 |
| Gene-PCA | SVM | 65.2 | **95.6** | 24.2 | 38.6 |
| All-PCA | LogisticRegression | 71.8 | 63.2 | **90.4** | 74.4 |
| All-PCA | SVM | 64.3 | 94.1 | 22.5 | 36.3 |
| - | Polygenic Risk Score* | 73.7 | 89.8 | 47.3 | 62.0 |

#### Solution

The solution is to apply linear techniques, such as PCA, only for small, biologically well defined functional units of the genome. The most obviously plausible such functional unit is a single gene.

Unlike *genome-scale* application, *gene-scale* application of linear techniques, such as PCA, preserves the possibility to detect meaningful, non-additive effects across genes. This is further supported by the argument that learning high level representations for variants in each gene individually preserves the local spatial structures of the genome, and by the fact that the majority of non-additive interactions are due to inter-gene regulatory effects, whereas within-gene non-additive patterns are less obvious [18, 47]. Further, genetic interaction networks form hierarchical structures that capture modules corresponding to protein complexes and pathways, biological processes, and cellular compartments [48]. Inter-gene interactions to establish phenotype that matter are commonly described as epistasis, canalization, robustness, or buffering [49], which, importantly, have been found to contribute to missing heritability as well [50].

Our experiments confirm these ideas, see Table 1: while the step from genome-scale (‘All-PCA’ in Table 1) to only gene-scale application of PCA (‘Gene-PCA’ in Table 1) considerably boosted the predictive power of the capsule network in use, employing non-additive dimensionality reduction techniques, such as autoencoders instead of PCA did not improve results any further (Supplementary Table 1). In fact, the application of non-linear reduction techniques rather yielded prediction runs to be unstable, and supported subsequent overfitting of the input data (Supplementary Table 1).

### DiseaseCapsule: Network Architecture

See Fig. 2 for an illustration of the following brief description of the architecture of DiseaseCapsule and Methods for full details. In the following, we focus on describing the network in relation to input resulting from the new, gene-scale dimensionality reduction protocol.

**Fig. 2.**
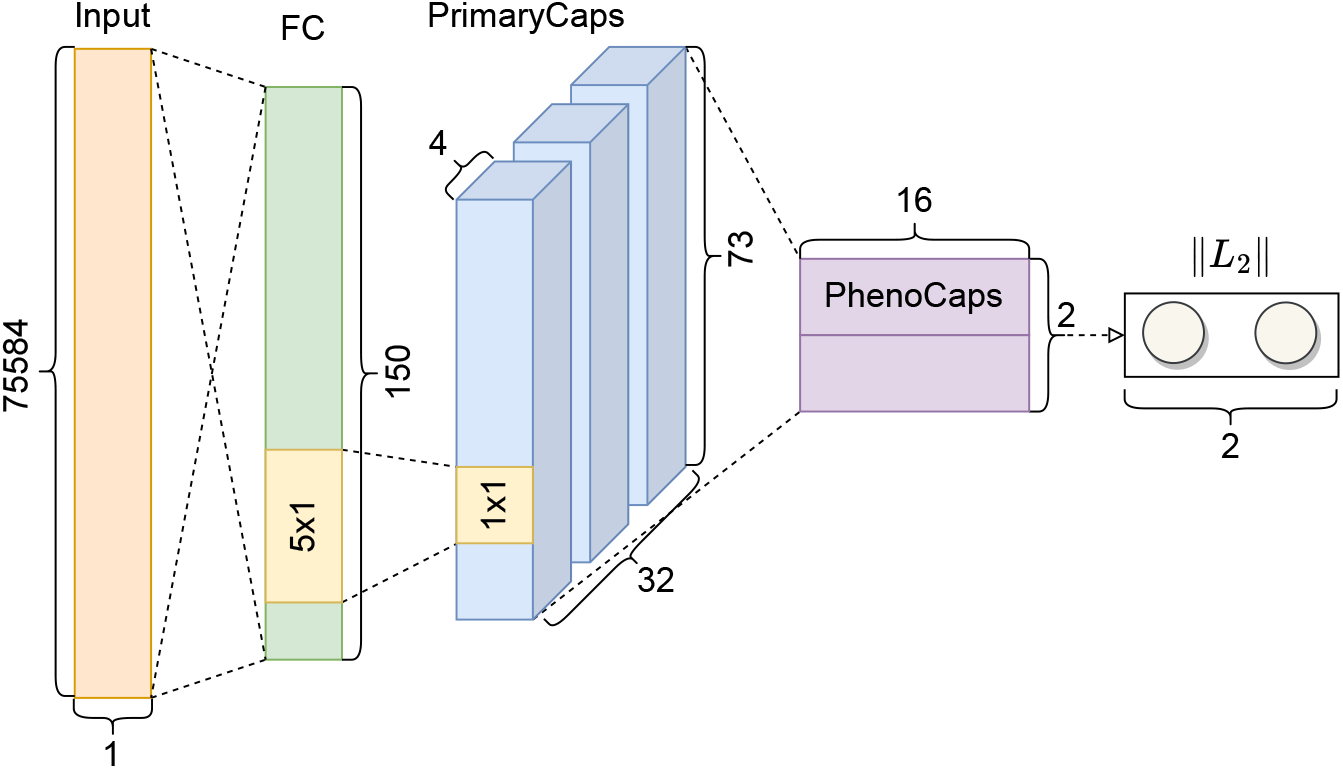
The architecture of DiseaseCapsule.The input is the concatenation of the compressed features from all Gene-PCA models, where each feature corresponds to one Gene-PCA. The number of Gene-PCA’s is 75584, so the dimensionality of the input is 75584×1. DiseaseCapsule consists of three layers: a Fully Connected Layer (FC), a Primary Capsule Layer (PrimaryCaps) and a Phenotype Capsule Layer (PhenoCaps). The FC layer consists of 150 neurons followed by ReLU as activation function. The PrimaryCaps is composed of 32 primary capsules. Each of them involves 4 different convolutional filters (kernel size = 5× 1, stride = 2, no padding). PhenoCaps consists of two 16-dimensional vectors. Each phenotype capsule receives input from all 32 primary capsules. The output is a binary classification label (Healthy or ALS).

The number of features arising from the reduction step amounts to 75,584. As explained before, each of the features refers to a linear combination of variants within a gene, where genes are defined according to the default annotation database in ANNOVAR [51], see Methods for details. This means that the input data is a vector of 75,584 real-valued entries for each patient. For a fair comparison, we reduce the features to the same number (75,584) when performing ‘All-PCA’ (i.e. standard, whole genome wide PCA).

This vector then is run through a fully connected layer (FC), followed by a primary capsule layer (PrimaryCaps) and a phenotype capsule layer (PhenoCaps). The combination of PrimaryCaps and PhenoCaps reflects, apart from minor adaptations that are necessary to capture the particular input data and the prediction task, a standard capsule network core architecture, as originally suggested [35]; note that unlike originally suggested, here capsule layers are preceded by a fully connected layer instead of a convolutional layer. This takes into account that the input is sequential in nature, and not a rectangle of pixels as in the original application.

The output of PhenoCaps is binary-valued, where the two values indicate prevalence or non-prevalence of ALS in the individual from which the input was retrieved. The prediction is thus straightforwardly established by evaluating the output of the PhenoCaps layer.

### Predicting ALS: Gene-PCA + DiseaseCapsule outperforms alternative approaches

See Table 1 for the following. We evaluated the performance of DiseaseCapsule with various state-of-the-art prediction approaches, such as multilayer perceptrons (MLP), logistic regression, support vector machines (SVM), a standard convolutional neural network (CNN) whose architecture resembles that of DiseaseCapsule, but uses convolution instead of capsule based layers, as well as Random Forests and AdaBoost. This represents a selection of predominant, high-performance machine learning approaches. Last but not least, we computed predictions based on polygenic risk scores (PRS) relative to different, reasonable GWAS thresholds. Note that computation of PRS does not require data reduction other than selecting SNP’s according to the different GWAS thresholds, which crucially distinguishes it from ’Logistic Regression’.

To further evaluate the contribution of the novel gene-scale dimensionality reduction protocol, we combined each method with standard genome-scale PCA (‘All-PCA’) or the novel protocol (‘Gene-PCA’).

The first observation is that all neural network based (so fully non-additive) models profit from using ‘Gene-PCA’ instead of ‘All-PCA’: accuracy of DiseaseCapsule, MLP and CNN increase from 81.9 to 86.9, from 72.5 to 84.2 and from 53.8 to 74.5, respectively. All other approaches can be run on ‘Gene-PCA’ or ‘All-PCA’ at near-identical performance (Random Forest and AdaBoost even incur losses in terms of accuracy when applying ‘Gene-PCA’). This points out that Gene-PCA predominantly caters to neural network models. From a conceptual point of view, this was to be anticipated, because Gene-PCA preserves the potential to detect non-linearities across genes, whereas ordinary PCA does not. Since applying local linearization prior to a linear model based analysis still results in an overall linear approach, as the concatenation of two linear functions, linear methods cannot profit from Gene-PCA; still they fail to pick up non-linearities. Note that although Random Forest or AdaBoost are not additive, the ’universal approximation theorem’ does not apply for them. This may be the likely explanation for why Random Forest and AdaBoost do not profit from Gene-PCA as much as DiseaseCapsule itself does.

In an overall account, DiseaseCapsule achieves prediction accuracy of 86.9, which establishes the top performance, rivaled only by MLP, as an alternative non-additive approach (84.2). The third best performance is achieved by DiseaseCapsule on ‘All-PCA’ (81.9), closely followed by PRS at a GWAS threshold of 5 × 10^-2^ (81.8). All other performance rates drop below 79. This means in particular that in a relative comparison with PRS, as a standard prediction technique, DiseaseCapsule leaves 28% less individuals misclassified, which establishes considerable, relevant progress, both from the point of view of clinical applications and from the point of view of predictive power in general. It also establishes a first quantification of the contribution of non-additive constellations of variants / genes to identifying ALS (see below for further analyses).

Again returning to DiseaseCapsule correctly classifying 86.9% of individuals one realizes that DiseaseCapsule is able to recognize genetic patterns that do or do not characterize the disease in 86.9% of the tested individuals. In other words, when considering DiseaseCapsule as a piece of artificial intelligence, DiseaseCapsule ’understands’ the patterns that drive the disease or not, in 86.9% of the individuals.

We then dove further into evaluating how many affected people were revealed by DiseaseCapsule as affected (as summarized by ’Recall’), and how many people not affected were misclassified (as can be measured by calculating ’Precision’). Both of these statistics are crucial measures for application purposes: while Recall indicates how many people affected are identified, so can be suggested for therapies, Precision indicates how many people are mistakenly suggested for therapies, hence are likely to suffer from possible side effects. Both cases should be prevented.

As a result, DiseaseCapsule clearly reveals itself as the most balanced and overall decidedly strongest approach. DiseaseCapsule achieves Precision of 85.2 and Recall of 89.4, which combines into an F1-Score of 87.2. This clearly outperforms the second best approach (again Gene-PCA + MLP), which reaches an F1-score of 82.6, where for MLP Precision (92.2) dominates Recall (74.8). That is, Gene-PCA + MLP overlooks more than a quarter of the ALS cases. Remarkably, All-PCA + Logistic Regression, coming out at an F1-Score of 81.4, reveals itself as the most aggressive approach: 96.0% of the cases are recognized as such; this however is offset by poor Precision: the approach incorrectly classifies 29.3% of healthy individuals as ALS cases. While the aggressiveness of the linear approach is remarkable, the many errors in labeling individuals as ALS patients put this into perspective: too many healthy individuals are sent to therapy. Again, PRS, by using a threshold of 5 × 10^-2^, reaches an F1-Score of 79.4.

Note finally that all approaches other than DiseaseCapsule, MLP or Logistic Regression / PRS achieve an F1-Score of only at most about 70. The only noteworthy phenomenon is the high Precision of SVM’s, amounting to 94.8 both for Gene- and All-PCA. However, Recall remains at a sobering 55.8 (again both for Gene- and All-PCA), meaning that SVM’s does not recognize nearly half (44.2%) of the ALS cases.

As for comparing different thresholds when making use of PRS, the least stringent threshold of 5 × 10^-2^ clearly outperforms the more stringent thresholds of 5 × 10^-4^, 5 × 10^-6^ and 5 × 10^-8^ (where performance steadily decreases on increasing stringency). At the same time, however, PRS at 5 × 10^-2^ is by far the most demanding approach in terms of memory requirements: it claims 160 GB of RAM while running, which is easily explained by the overwhelming amount of variants that need to be processed.

See further Supplementary Table 2 for results referring to the methods Bayesian LASSO (BL), Genomic Best Linear Unbiased Predictor (GBLUP), Bayesian Ridge Regression (BRR) and BayesB, all of which are common for predicting phenotype from genotype. All these methods obtain less than 70% accuracy and, as a particular potential drawback, less than 45% recall. This points out that these methods are rather inferior with respect to the approaches presented in Table 1.

### Validating DiseaseCapsule on Parkinson’s Disease

To validate the predictive performance of DiseaseCapsule also in other diseases with a genetic background, we applied DiseaseCapsule as well as the other representative models on Parkinson’s disease (PD) data [52, 53, 54, 55], following the exact same protocol as for ALS. See Supplementary Table 3 for the results achieved on the corresponding test set. The results show that performances of all approaches in PD amount to not more than 62% accuracy (DiseaseCapsule). In a summary of results, in analogy to ALS, Gene-PCA + DiseaseCapsule outperforms all other approaches in terms of accuracy (62.0%), recall (68.1%) and F1 score (64.2%) in PD.

Based on the design of DiseaseCapsule and based on heritability estimates that one can obtain from the literature, we reckon that the differences in terms of performance between ALS on the one hand, and Parkinson’s on the other hand, have two explanations. First, the considerably reduced (> 30 times lower) number of SNP’s being captured by the Parkinson’s data appears to put limitations on the accuracy in prediction that one can achieve. Note that the DiseaseCapsule protocol is tailored towards being able to handle large amounts of variants, also because DiseaseCapsule is supposed to support the omnigenic model; apparently, smaller amounts of SNP’s have an impact on all methods, including DiseaseCapsule. Second, heritability estimates raised in the literature for Parkinson’s expose Parkinson’s to be considerably less affected by missing heritability [12, 13, 54, 56]. This is a hint towards non-additive interactions between variants and genes playing a less important role than for ALS.

### Classification performance improves on increasing numbers of genes

We recall that PRS models steadily improved by integrating increasing numbers of genes, obtained by relaxing GWAS significance thresholds. Similar to this finding, we were interested in how DiseaseCapsule performed relative to decreasing or increasing numbers of genes. To do that, we randomly selected a certain number of genes, and set all entries in the input vector (each of which refers to one within-gene principal component) that referred to non-selected genes to zero. We then ran the resulting, modified vectors of all 1040 test samples, and evaluated the runs in terms of Precision, Recall, F1-Score and Accuracy. We did this for each of a given number of genes, where numbers increased from 1, 50, 100, 200, 500, 1000, 2000, and further in steps of 2000 until 18,000 (i.e. nearly the overall number of genes). For each given number of genes, we repeated the random selection process 300 times, and evaluated each run in terms of Precision, Recall, F1-Score and Accuracy. We arranged the corresponding results as boxplots with each box referring to the 300 runs for one given number of genes. See Fig. 3 for the corresponding results.

**Fig. 3.**
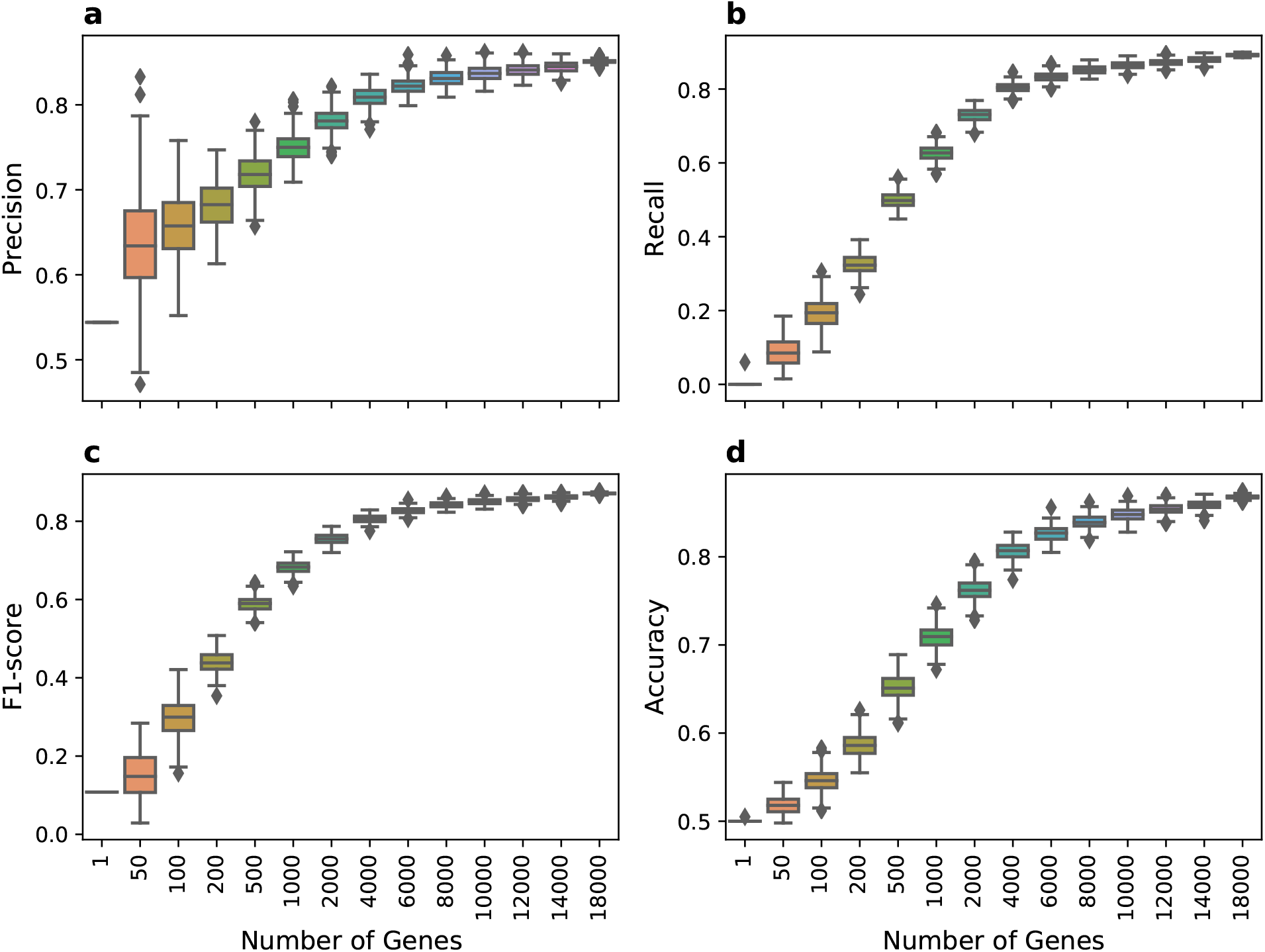
Prediction performance from randomly chosen genes. Precision, Recall, F1-score and Accuracy are shown in panel **a**, **b**, **c**, **d**, respectively. The evaluation process was repeated for 300 times for a specific number of selected genes. The x-ticks in four figures are the same.

It becomes immediately evident that increasing numbers of genes improve results in all aspects, which, to some extent, can be explained by the omnigenic model of diseases [14]. Of note, increases in performance are dramatic up to 4000 genes, while from 6000 genes onwards, improvements are rather minor.

Intriguingly, the variance in terms of performance (see e.g. Fig. 3d) can be very large when using at most 2000 genes, but drops substantially from at the latest 6000 genes and more. This means that outstanding performance can be achieved on only 2000 genes already, if these have been selected preferentially.

It remains to design a strategy through which to select the genes that are most relevant for classification; most likely, such genes have key roles in establishing or preventing the disease.

### Determining genes decisive for classification

See Fig. 4 and Online Methods for explanations in the following. In the DiseaseCapsule network, the vectors of the two phenotype capsules ‘ALS’ and ‘Healthy’ (**s_j_** in Fig. 4), consist of linear combinations of output vectors provided by the primary capsules (**u_j|i_** in Fig. 4). The linear weights *c_ij_* that connect the **s_j_** with the **u_j|i_** are referred to as coupling coefficients. Unlike ordinary parameters of the network (e.g. **W_ij_** in Fig. 4), the *c_ij_* are determined through the dynamic routing procedure, which does not make part of core training through backpropagation. As per its nature, the dynamic routing algorithm is such that a situation with little—usually even only one—large *c_ij_* is favored over several equally large *c_ij_* for each *j*.

**Fig. 4.**
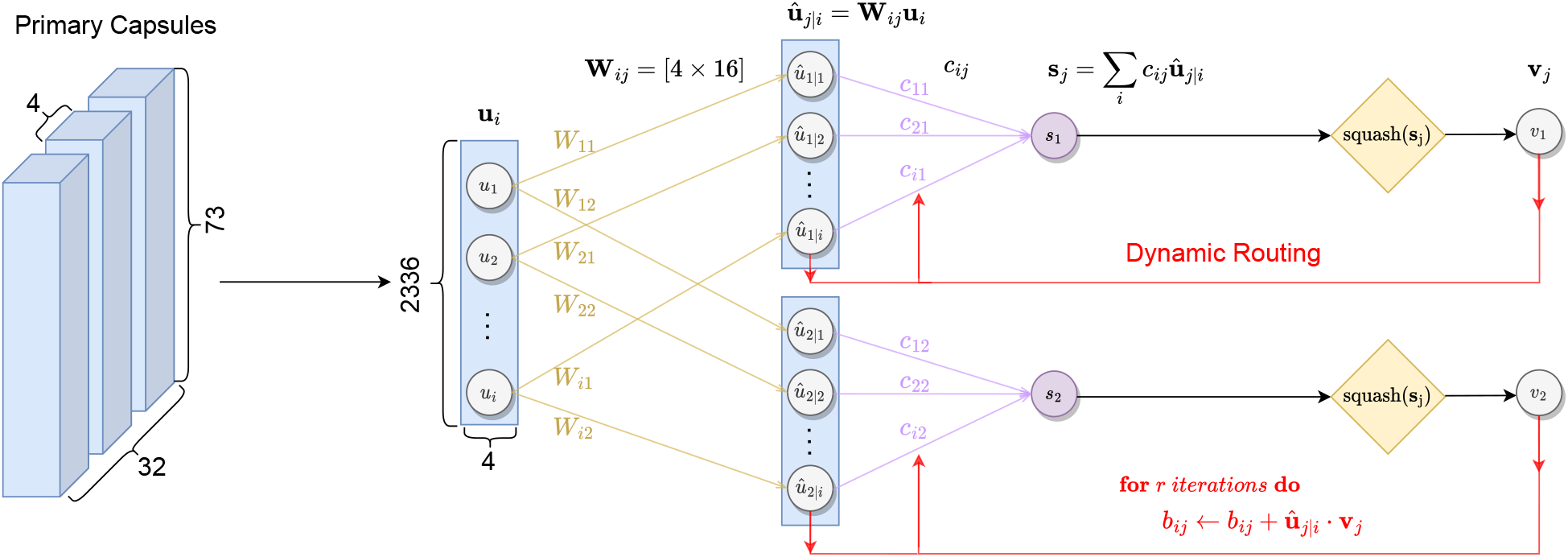
The schematic diagram of the dynamic routing algorithm. The PrimaryCaps layer outputs 32 × 73 = 2336 vectors **u**_*i*_(*i* = 1,…, 2336), which are then transformed into 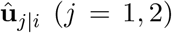 by premultiplying pose matrices **W**_*ij*_. Note that pose matrices are learnt during training. The cycles starting with purple arrows and ending with red arrows show the iterative dynamic routing procedures. The coupling coefficients *c_ij_* indicate the probability that primary capsule *i* agrees with phenotype capsule *j*. The input **s**_*j*_ to one of the phenotype capsules is subsequently processed by using the squashing operation. The output **v**_*j*_ is used to update the *b_ij_* and *c_ij_* until the model converges.

Further, primary capsules that have great coupling coefficients in connection with the ‘ALS’ capsule, are likely to code for constellations of genes that drive the disease. Vice versa, primary capsules sharing links with the ‘Healthy’ output capsule that are equipped with large coupling coefficients, code for genes whose activation (or de-activation) distinguishes healthy individuals from the ones affected with ALS. Importantly, as outlined above, the number of such primary capsules is (very) small.

Combining these ideas means that commonly very little or even only one primary capsule captures the decisive relationships among genes that distinguish diseased from healthy people. We therefore investigated how primary capsules related to the ‘ALS’ output capsule (which predominantly fires when ALS is to be predicted) on the one hand, and the ‘Healthy’ output capsule (which predominantly fires when the individual is to be classified as not being affected by ALS) on the other hand.

To highlight primary capsules either being important for activating the ‘ALS’ or the ‘Healthy’ output capsule (as documented by large coupling coefficients on the respective primary-output capsule links), we ran all training and all test samples through the network, amounting to two separate runs, one for the training and one for the test data. The intuition is to demonstrate that despite not having been part of the training, effects reproduce on data that had not been seen before. We collected the resulting coupling coefficients; note that coupling coefficients are computed individually for each sample by way of the ’dynamic routing’ protocol during the forward pass [35]. So, coupling coefficients are not determined as part of backpropagation during training, which would imply that coupling coefficients are the same for all individuals. We averaged the resulting coupling efficients across all samples, for each of the 2 × 32 possible combinations of PrimaryCaps and PhenoCaps. As above-mentioned, we did this for both training and test data. The corresponding averaged coupling coefficients are displayed in the two heatmaps in Fig. 5, where Fig. 5a is for the training data and Fig. 5b is for the test data. See also Methods for further details on the experiments and the corresponding visualization process.

**Fig. 5.**
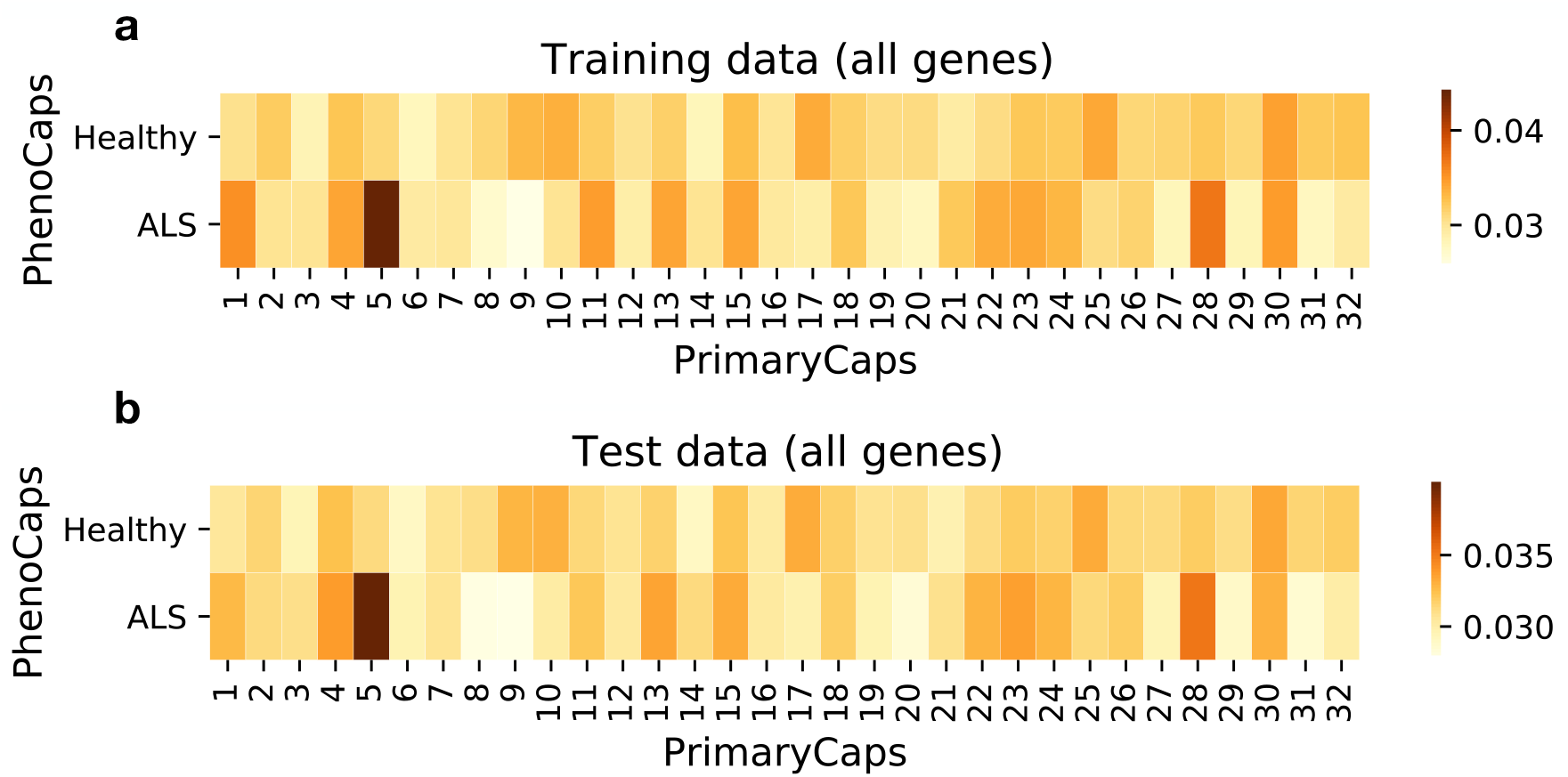
Heatmap plots for average coupling coefficient matrices of DiseaseCapsule. **a**, Only plot for training individuals. **b**, Only plot for test individuals. The x-axis and y-axis represent the 32 primary capsule groups and 2 phenotype capsules, respectively. All 18,279 genes involved in DiseaseCapsule model are retained when predicting on test samples.

The most striking effect is that primary capsule 5 establishes the strongest link to the ‘ALS’ output capsule, both for training and test data. Far lesser so, but still apparent, primary capsule 28 activates the ‘ALS’ capsule. The agreement between training and test data demonstrates that effects do not only get manifested on data that was used to establish the parameters of the network (namely, the training data). In summary, activation of primary capsule 5 is the by far predominant effect from which to determine whether an individual is affected with ALS or not.

Correspondingly, we developed an algorithmic protocol according to which to determine core genes that markedly contribute to the activation of primary capsule 5, see Methods for details. Using this protocol, we obtained 922 core genes that contribute to the activation of primary capsule 5 (Fig. 6a). This means that these 922 genes are important for DiseaseCapsule to identify the occurrence of ALS. To validate the predictive power of these 922 genes, we masked all other genes in the test data (i.e. we set entries in the input vector to zero if referring to genes not among the 922 selected ones), and ran the modified test data through the (trained) DiseaseCapsule model. As a result, using these 922 genes alone—which as we recall are crucial for activating primary capsule 5—a test accuracy of 76% was achieved. Note that random selections of 922 genes 70% accuracy on average, with a standard deviation of 1%, see Fig. 6b for corresponding results. So, the 76% achieved by the genes selected through our protocol are significantly greater (p-value < 2.2E-16).

**Fig. 6.**
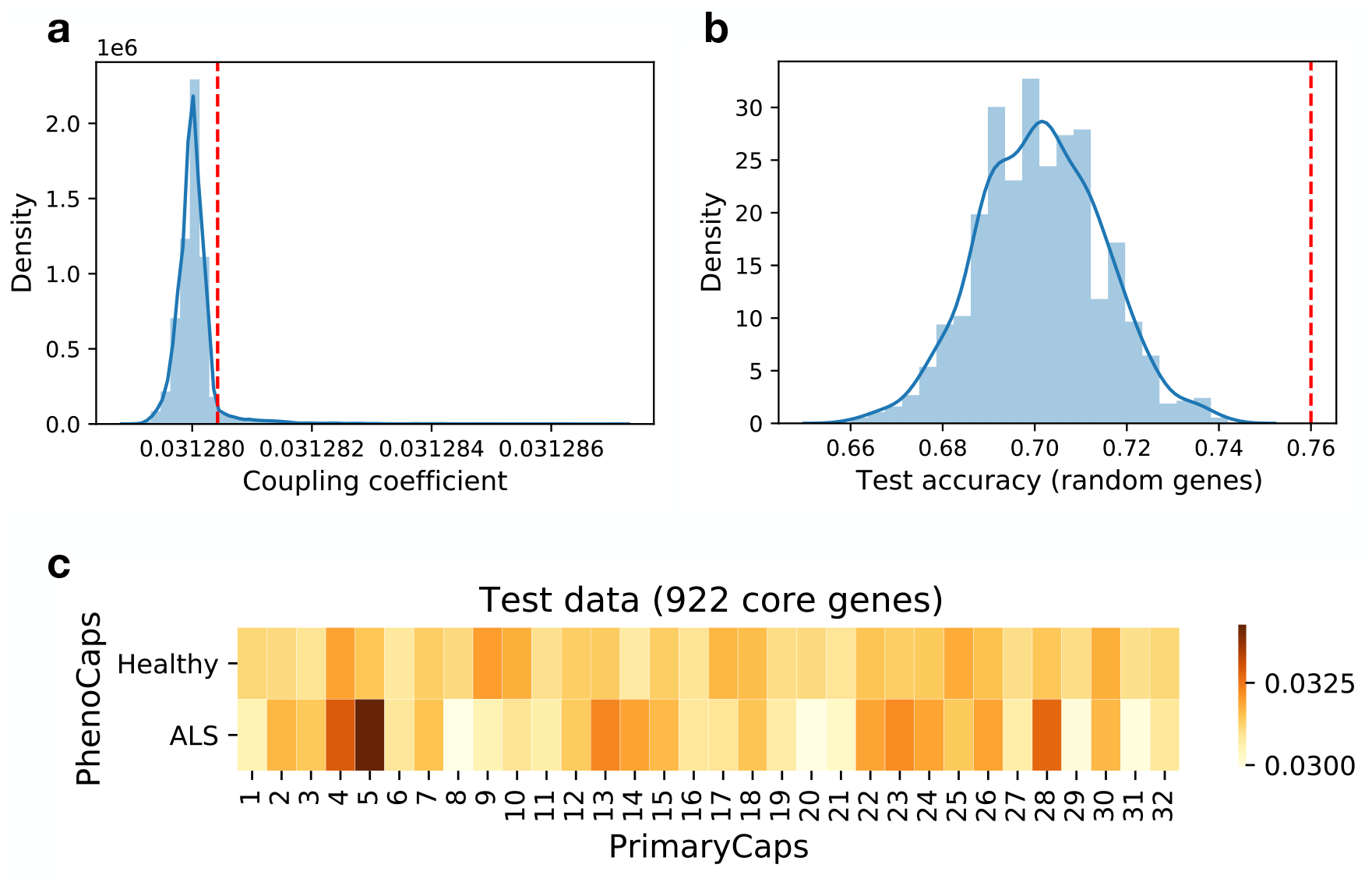
Determining and validating core genes decisive for classification. **a**, The distribution of coupling coefficients between primary capsule 5 and phenotype capsule ALS for all genes. The red dashed line indicates the 95-percentile. 922 genes whose coupling coefficients are above the 95-percentile are selected as core genes decisive for classification. **b**, Test accuracy distribution of using 922 randomly chosen genes as input for DiseaseCapsule model (repeat for 1000 times), while the other genes are masked (set as zero). The red dashed line indicates the test accuracy of using 922 core genes as input. **c**, Heatmap for average coupling coefficient matrices (test data) using 922 core genes as input.

To further corroborate the predictive power of the 922 selected genes, we computed the average of the coupling coefficients across all test individuals when using only the 922 preferentially selected core genes, see Fig. 6c. The level of activation of primary capsule 5 (0.0342) nearly matches the one when using all genes (0.0399).

### Examining and classifying 922 genes decisive for classification

It therefore made sense to examine and classify these 922 genes further; see Supplementary Table 4 for the resulting list.

When examining these 922 potential core genes according to their classification, we find that 13 genes *ARHGEF28, BCL11B, C9orf72, CDH13, CNTN4, CNTN6, DPP6, ERBB4, ITPR2, LIPC, RAMP3, SOX5, SYNE1* are also listed in the Amyotrophic Lateral Sclerosis online Database (ALSoD, https://alsod.ac.uk/), which so far includes 126 ALS-related genes. Note that *C9orf72* has been reported as the most common gene associated with ALS, accounting for about 6% of patients in white Europeans [57, 58]. Despite the few overlaps of our list with those in the database, we note that our genes agree with the ones in the database in terms of biological function (referring to neurological disorders in general) in various cases. In addition, 52 genes that we found also make part of the 690 ALS risk genes that have recently been revealed by exploiting non-linearities within LD blocks [22].

To quantify the overlap in terms of functional annotations more rigorously, we performed a Gene Ontology enrichment analysis for these 922 genes. The analysis shows that our potential core genes are significantly enriched in molecular function terms such as: calcium ion binding (GO:0005509, adjusted P value (Padj) = 9.6E-04), etc, and enriched in biological process terms such as: nervous system development (GO:0007399, Padj = 2.5E-13), neuron differentiation (GO:0030182, Padj = 4.0E-11), etc, and also enriched in cellular component terms such as: synapse (GO:0045202, Padj = 6.7E-11), plasma membrane (GO:0005886, Padj = 2.0E-09), and so on; see details in Supplementary Fig. 1.

An additional pathway enrichment analysis further demonstrated that these 922 genes are significantly enriched in pathways such as: protein-protein interactions at synapses (R-HSA-6794362, Padj = 1.2E-02), Netrin-1 signaling (R-HSA-373752, Padj = 1.3E-02), and also some more pathways meaningful in terms of the disease. See Supplementary Fig. 1 for details.

In summary, the collection of enriched GO terms and pathways have been shown to significantly relate with human nervous system related diseases, which do include ALS in most cases [59, 60, 61, 62, 63, 64]. Since the 922 genes act as decisive variables in the functional relationship by which one computes phenotype from genotype (namely, the neural network function reflected by DiseaseCapsule), they play a role in taking genotype to phenotype, from a merely functional point of view. In that sense, we believe that many of them could have the potential to act as genes that are involved in the genetic architecture of ALS.

### 644 “non-additive” genes

We used a simple target function that measures the difference between the contribution of a selection of genes to correct classification in DiseaseCapsule, and the contribution of a gene in a common logistic regression scheme, quantified in both cases by the accuracy that one achieves when running the two methods on the selected genes alone. Employing a genetic algorithm, we determined a subset of 644 genes that maximized the target function values. That is, running methods on these 644 genes maximizes the difference in the non-additive scheme (DiseaseCapsule) and the additive scheme (regression), see Methods for full details. In other words, these 644 genes are crucial for classification in non-additive schemes, but remain useless when being used in a linear scheme, because the linear scheme is blind to their effects; more than that, these 644 genes reflect a selection that is optimal in that respect.

To reinforce that these 644 “non-additive” genes are responsible for predominantly non-additive effects, we ran both DiseaseCapsule (Gene-PCA + DiseaseCapsule in Table 1) and Logistic Regression (Gene-PCA + Logistic Regression, which proved the strongest additive protocol, see Table 1), both on training (which led to establishing parameters for both DiseaseCapsule and LogisticRegression) and test data (hitherto unseen by both approaches). The differences in accuracy are striking, both on training (difference: 0.227) and test (difference: 0.162) data.

Although striking, the difference between training and test data may be due to hidden biases. To ensure that these 622 genes nearly exclusively yield non-additive effects—which can only be picked up by DiseaseCapsule—we retrained both DiseaseCapsule and Logistic Regression on these 622 genes alone. This gives both approaches the same fair chance: try to get the most out of the 622 genes they have been provided with.

See Table 2 for the corresponding results. Retraining both models yields accuracies of 0.712 for DiseaseCapsule, but only 0.512 for Logistic Regression (amounting to a difference of exactly 0.2, matching the range of the earlier results). Note that because test data was balanced (equal cases and controls), 0.5 means random performance. So, on these 622 genes, Logistic Regression matches random classifier rates, while DiseaseCapsule achieves excellent performance rates.

Breaking down results in terms of Precision and Recall confirms that DiseaseCapsule establishes decent performance rates (Pr: 0.715; Rec: 0.706), whereas Logistic Regression’s performance does not even match minimum standards (Pr: 0.522; Rec: 0.292(!)).

#### Functional annotations of the 644 non-additive genes

The corresponding 644 “non-additive” genes are listed in the Supplementary Table 5. We ran a brief analysis on the 644 non-additive genes using Reactome [65]. The corresponding pathway enrichment analysis demonstrates that some genes are significantly enriched in pathways that are related with neurodegenerative diseases [66, 67, 68]. Examples of such pathways are “MECP2 regulates transcription of genes involved in GABA signaling” (pValue=2.54E-4), or “HSF1-dependent transactivation (pValue=8.78E-4)”.

An additional observation of interest is that 20 of the 644 genes had been revealed by the recent study that took local (but not global) non-linearities into account [22]; importantly, *KANK1*, which was particularly highlighted in [22] and confiremd by convergent genetic and experimental data, also belongs to the 644 non-additive genes presented here.

### DiseaseCapsule is less sensitive to batch induced / cohort specific confounding effects

The data set we used is composed of four batches, with technical identifiers C1, C3, C5 and C44. The number of cases and controls in the large batches C5 and C44 is highly unbalanced. While C5 consists of controls alone, C44 consists of predominantly cases (80% cases). See Supplementary Table 6 for details. The obvious concern is that a classifier picks up batch structure (so, for example, predominantly distinguishes C5 from C44), instead of really distinguishing controls from cases. To alleviate this problem, we removed all SNP’s prone to batch effects based on a statistical evaluation prior to main analysis (see Subsection “Data: Project MinE”).

Nevertheless, some less obvious batch effects may have remained despite this explicit correction. We therefore evaluated the performance of all classifiers that yielded above average performance in the main analysis (DiseaseCapsule, MLP, CNN, Logistic Regression, SVM, PRS). To not be able to pick up confounding batch effects due to the dominance of the heavily unbiased batches C5 and C44, we used C5 and C44 as training data (amounting to 92.5% of the samples overall), an evaluated the leading methods on C1 and C3 as test data (amounting to 7.5% of the samples). Since biases due to batch structure in C5 and C44 cannot reproduce in independent batches, the arrangement ensures an evaluation free of batch induced biases. Last but not least note that numbers of cases and controls is roughly balanced in C1 and C3 (355 cases vs. 429 controls together), so warrants a sufficiently fair evaluation.

The corresponding results are shown in Table 3. Gene-PCA + DiseaseCapsule still achieves the best accuracy (81.6%) and F1 score on the test data (C1 & C3). While this means a loss in performance of 5.3%, other approaches suffer considerably more. Gene-PCA + MLP, All-PCA + Logistic Regression, Gene-PCA + SVM and PRS get worse by 9.7%, 6.3%, 11.1% and 8.1%, respectively. In addition, while Gene-PCA + DiseaseCapsule can preserve high precision (85.8%) at still good recall (71.3%), other approaches incur significant losses in particular in terms of recall, dropping below 50% in all cases but Logistic Regression (90.4%, at Precision 63.2%). Of note, MLP and SVM both achieve excellent precision (94.3% and 95.6%, at Recall 46.5% and 24.2%, however).

Overall, Gene-PCA + DiseaseCapsule outperforms other approaches even more clearly (both Accuracy and F1-Score are substantially larger than those of alternative approaches) than on the basic, non-batch adapted data. This points out that Gene-PCA + DiseaseCapsule is not only the best approach in terms of basic performance. It is also the approach that apparently resists batch induced biases clearly the most.

### DiseaseCapsule needs less data for training

Standard deep neural networks, such as CNN’s tend to require massive amounts of training data for achieving optimal performance. In the seminal paper [35] it was pointed out that capsule networks require substantially less data for training than ordinary CNN’s. This can be attributed to the “viewpoint invariance property”, which, as above-mentioned, means that capsule networks can recognize elements in images (such as houses or cars) from various angles, although training data from only one particular angle was provided. We recall that training ordinary CNN’s do not enjoy this property. This means that training data referring to various angles need to be provided in order to have CNN’s recognize objects from various angles.

To evaluate whether corresponding effects persist also for the genotype profile data under consideration here, we reduced the training-validation dataset by subsampling it, reducing it in size from 100% to 20% in steps of 20%, and further 10% and 5%, repeating the subampling 10 times to account for random fluctuations. In case of 100%, we randomly initialized training runs 10 times on the identical (full) data, so as to account for baseline fluctuations due to initialization. We then trained all models on only the subsampled data (using identical hyperparameters in all cases). Results are shown in Table 1. This process was repeated for ten times. All models were evaluated on the same test dataset.

Results shown in Table 4. Obviously, DiseaseCapsule outperforms all other approaches relative to all subsampling rates. Further, the first subsampling rate at which approaches loose more than 5% varies. While DiseaseCapsule reaches this point only at 20% (86.3 → 81.3), MLP, SVM, CNN and PRS already cross the 5% line of losses earlier: 40%, 60%, 40% and 40%, respectively. The only alternative approach that proves stable with respect to reductions in data is basic Logistic Regression, whose performance drops by 5.2% at a subsampling rate of 20%, just as DiseaseCapsule.

**Table 4.** Test accuracy of various models trained using different percentage of subsampling training samples, namely 100%, 80%, 60%, 40%, 20%, 10%, 5%. The first column denotes the subsampling percentages. Values are represented as mean ± sd (%), which are calculated for the same test dataset by repeating the subsampling and training-testing processes for ten times with the identical network architecture and hyperparameters. DR: dimensionality reduction, LR: Logistic Regression. PRS: Polygenic Risk Score. *This model is PRS-based that the SNPs were selected by GWAS with the threshold (*P* < 5 × 10^-2^). *Note*: The best value is marked in bold.

| DR | Gene-PCA | Gene-PCA | Gene-PCA | Gene-PCA | Gene-PCA | - |
| --- | --- | --- | --- | --- | --- | --- |
| Classifiers | DiseaseCapsule | MLP | LR | SVM | CNN | PRS* |
| 5% | **74.0** ± 1.4 | 61.5 ± 1.1 | 69.1 ± 0.5 | 50.0 ± 0.0 | 53.7 ± 2.0 | 60.1 ± 1.1 |
| 10% | **79.0** ± 0.8 | 66.0 ± 0.9 | 72.1 ± 0.5 | 53.2 ± 0.3 | 56.2 ± 1.6 | 67.0 ± 0.8 |
| 20% | **81.3** ± 0.6 | 71.4 ± 0.6 | 73.4 ± 0.3 | 61.4 ± 0.2 | 61.7 ± 2.0 | 71.9 ± 0.8 |
| 40% | **83.5** ± 0.5 | 76.5 ± 0.6 | 74.6 ± 0.4 | 67.3 ± 0.3 | 66.8 ± 0.9 | 76.9 ± 0.9 |
| 60% | **84.2** ± 0.3 | 79.9 ± 0.7 | 76.1 ± 0.3 | 71.1 ± 0.1 | 70.2 ± 1.3 | 79.3 ± 0.6 |
| 80% | **85.1** ± 0.4 | 81.5 ± 0.6 | 76.7 ± 0.3 | 73.2 ± 0.2 | 72.3 ± 1.1 | 80.8 ± 0.5 |
| 100% | **86.3** ± 0.3 | 83.6 ± 0.9 | 78.6 ± 0.3 | 76.1 ± 0.2 | 73.7 ± 0.7 | 81.9 ± 0.3 |

In summary, relative to the optimal performance, only DiseaseCapsule and Logistic Regression prove to be able to work with relatively little data, without incurring significant losses. In that respect, importantly, DiseaseCapsule is the only more sophisticated machine learning approach that can work with little data.

Last, please see Supplementary Table 7 for a corresponding evaluation in terms of Recall and Precision. Also these results prove DiseaseCapsule and Logistic Regression react most favorably with respect to reductions in training data.

Moreover, DiseaseCapsule keeps good performance in terms of precision and recall when subsampling training data (Supplementary Table 7). Notably, DiseaseCapsule obtains 81.7 ± 1.9% precision and 81.7 ± 2.8% recall when subsamping 20% training samples. However, the recall of other methods except Logistic Regression goes down drastically, none of which can exceed 50%. For example, MLP, CNN, SVM and PRS-based model can only obtain 46.7 ± 1.5%, 36.2 ± 6.8%, 22.9 ± 1.5%, 48.5 ± 1.0% in terms of recall, respectively. Though when subsampling 20% training data All-PCA + Logistic Regression still achieves the best recall (85.8 ± 1.6%), it performs worse in terms of precision (73.7 ± 1.3%) and accuracy (77.6 ± 1.4%) compared with DiseaseCapsule model. Additionally, DiseaseCapsule always shows much better performance compared with CNN in terms of accuracy, recall and sensitivity to resampling and subsampling. In general, our DiseaseCapsule model is robust and shows good performance when training with less data, which outperforms other approaches in most cases.

### Runtime and memory usage of our approach are acceptable

Gene-PCA was run on a CPU cluster. Running Gene-PCA on a single gene of 100 SNP’s took approximately 17 seconds at peak memory of 0.25GB. Note that the Gene-PCA step can be run in parallel on a multi-core machine. Further, DiseaseCapsule required 14 minutes and 9GB of RAM for training when running on a GPU (Nvidia GeForce RTX 2080 Ti).

## Discussion

In this study, we have presented DiseaseCapsule, as a novel deep learning based approach to infer disease phenotype from individual genotype. As the major novelty, *DiseaseCapsule captures non-linear, arbitrarily complex functional relationships* between genotype and phenotype *across the whole genome*. All prior approaches presented so far considered to evaluate variants in additive schemes (which reflects the common standard), presented approaches that consider non-additivity only within small local boundaries, or resorted to considering only a few genes, or some selected chromosomes when operating in more global ways.

DiseaseCapsule has come with two immediate theoretical advantages. First, because of operating across the whole genome, DiseaseCapsule does not need to focus on a few core disease-related genes only, so does not miss the weak effects of abundant peripheral genes. Second, DiseaseCapsule improves on capturing the hierarchical structures of the underlying genetic interactions thanks to the high degree of complexity that capsule networks can capture. This plays a particularly relevant role for ALS, because ALS is commonly hypothesized to have a complex genetic architecture.

In practical terms, DiseaseCapsule has outperformed all state of the art approaches when predicting amyotrophic lateral sclerosis (ALS) from individual genotype: it has achieved 87% accuracy on held out test data. This translates into a relative increase of 28% over polygenic risk scores (PRS), so remains with 28% less misclassified people in comparison with the current clinical standard. This establishes substantial, arguably even drastic advantages in comparison with what was possible earlier. Analyzing results further has revealed that DiseaseCapsule achieves 89.4% recall, which reflects that DiseaseCapsule identifies 64% more ALS patients than the clinical standard (PRS: 70.2%). This certainly means a potential strength of DiseaseCapsule when applied in clinical practice.

DiseaseCapsule has also redeemed its two major theoretical promises for application in clinical practice: sustainable use of training input, which reduces costs and efforts when raising clinical data, as well as advances in terms of interpretability of predictions. The latter point has become obvious through experiments based on inspecting the individual capsules of the network. The experiments have revealed 922 genes likely associated with ALS, many of which had not been pointed out before, but all of which appear to be plausible according to their functional annotations.

Realizing these advantages has required to overcome various non-negligible technical hurdles. First, integrating whole genome data means dealing with feature spaces whose dimensions are in the millions, as the amount of polymorphic sites in the human genome. Here, we have developed a protocol that exploits the major advantages of principal component analysis (PCA), without disrupting the possibly non-linear interactions between genes (which standard, whole-genome application of PCA does). Note further that non-linear approaches to whole genome wide dimensionality reduction, such as autoencoders, resulted in overfitting the data and unstable training runs. The protocol we have developed yields gene-specific principal components, which can be applied also for highlighting non-linear interactions across genes. Secondly, capsule networks had never been applied to whole-genome genotype data before. We have enabled this by means of an architecture that uses fully connected, instead of convolutional layers. This preserves to capture interactions between genes across the whole genome to a maximal degree, and appropriately accommodates the sequential nature of the input.

While determining the exact reasons for the superiority of DiseaseCapsule in predicting ALS from genotype still requires further investigation, some plausible hypotheses can be raised already.

First, as already alluded to above, capsule networks have the potential to learn the intrinsic hierarchical structures that underlie the data: their capsules, as complex (vector-valued) entities capture such structures much better than single-valued neurons, as their counterparts in ordinary neural networks. The enhanced capability to analyze complex biological relationships that underlie diseases (see also [37]) explains the improved generalization over other models.

Second, as above-mentioned, ALS had been commonly described as a disease that has a considerable genetic background. However, despite the consensus in this agreement, ALS still suffers from a non-negligible amount of missing heritability. Because the major resource for revealing heritability are linear model based genome-wide association studies (GWAS), missing heritability is often ascribed to non-linear genetic interactions between variants (e.g. epistasis) that have remained overlooked. Unlike the standard approaches, DiseaseCapsule is able to pick up such effects. To further investigate this hypothesis, we considered an objective function that addressed to find genes that served classification in (the non-linear) DiseaseCapsule, but not in linear regression type schemes. Optimizing according to this objective function (as per a genetic algorithm) yielded a subset of 644 genes that significantly contributed to predicting ALS in DiseaseCapsule (accuracy: 71.2%), while not working within the frame of logistic regression (accuracy: 51.2%). Evidently, linear approaches remain blind to these 644 genes. Although these 644 genes still require further inspection, it is reasonable to assume that various genes among them have the potential to be of future use in exploring the heritability of ALS further.

Third, one can hypothesize that ALS follows an omnigenic model [14] to a non-negligible degree. Results for polygenic risk scores (PRS) corroborate this idea. Gradually relaxing the threshold for inclusion of variants from 5 × 10^-8^ to 5 × 10^-2^ increases the accuracy by 20%. This means that numerous variants of small effect contribute to establishing ALS, in addition to the core disease-causing genes. DiseaseCapsule follows a whole-genome approach that does not put significance thresholds on individual variants (or genes) to appropriately take this into account.

As already alluded to above, DiseaseCapsule requires less training data than other approaches to establish excellent performance. While the effects are obvious, the translation of the “viewpoint invariance property” into the realm of genes and diseases still needs to be provided. It is reasonable to hypothesize that capsule networks capture the core effects regardless of their distribution across the ancestors of the individuals. Instead, other approaches may become confused and require to be presented with all possible combinations of effects when combining haplotypes

Despite the very promising results presented, further promising avenues of research are conceivable. Immediate such ideas are to also integrate epigenetic information, for example, or moving from genotype to haplotype data, whenever available.

In addition, while having established a deep learning based approach to predict disease phenotype from genotype, it appears sensible to develop a formal protocol for machine learning based approaches by which to identify (combinations of) variants that are associated with the diseases / phenotypes one examines. While such formal association schemes do not yet exist, we have made some great steps towards that goal, by making use of capsule networks. Unlike conventional deep learning models, capsule networks do not suffer from the notorious complaint to reflect inscrutable black boxes. We recall that DiseaseCapsule has already been able to deliver 922 genes that have been crucial for the classification process, which is a necessary condition for being associated with the disease. Another potentially fruitful undertaking, of course, is to study (some of) the 922 genes regardless of the existence of a formal protocol, just because many of them can shed light on the complex interactions that underlie ALS already now.

Fortunately, the field of explainable deep neural networks is moving fast ([69] looks like a particularly promising approach for our purposes, for example). Thus, one can expect rapid progress in the nearer future. Application of such strategies will help to understand the molecular mechanisms that underlie diseases, and aid practitioners in their assessment of strategies for preventing and treating the diseases.

## Materials and Methods

### Data: Project MinE

The data we used is from Project MinE https://www.projectmine.com, a large-scale study that aims to reveal the epi-/genetic mechanisms that underlie ALS, in the frame of globally concerted collaboration [70]. The data we use in this project is from the Dutch cohort of the project. It contains 7213 healthy (a.k.a. control) individuals and 3192 individuals affected with ALS. The cohort counts 5208 females and 5197 males. All participants of the study were genotyped using Illumina 2.5M SNP array.

#### Quality Control

First, we annotated SNPs according to dbSNP137 and mapped them to hg19 as the reference genome. We first performed quality control (QC) so as to remove low quality SNPs and individuals of low quality overall, by using PLINK 1.9 [71, 72] (–geno 0.1 and –mind 0.1). We stratified individuals according to the genotyping platform, and subsequently performed a more stringent SNP QC (-geno 0.0, –maf 0.01, –hwe 1e–5 midp include-nonctrl). We kept only SNPs in autosomal regions, and filtered based on differential missingness (–test-missing midp), excluding SNP’s of a p-value above 1e–4. As a result, we obtained 4,370,685 SNPs all of which are contained in the intersection the four batches (see below) the data consists of (see Supplementary Fig. 2).

#### Batch Structure and Batch Effects

The data set consists of four batches, pertaining to technical identifiers C1, C3, C5 and C44. Numbers of samples and ratio of cases vs. controls can vary substantially across batches: C1 contains 225 cases and 380 controls, C3 is 130 cases and 49 controls, C44 no cases, but 5155 controls, whereas C44 finally contains 2387 cases and 1629 controls (Supplementary Table 6). It is important to realize that C5 and C44, both of which are highly imbalanced—C5 contains no cases, while the majority of C44 are cases—cover approximately 92.5% of the samples, so dominate the data set.

This points out that removing batch effects is important. Otherwise, predicting cases and controls can be confounded with predicting C44 from C5, at still decent performance rates.

To remove artefacts by which to distinguish C44 from C5, we considered only the 5155 and 1629 healthy individuals from C5 and C44, respectively. Any SNP that supports to discriminate between C5 and C44 healthy individuals can be identified as signaling batch induced effects, so should be removed from further analysis.

To filter for batch effect transporting SNP’s, we computed a 2 × 2 contingency table for each SNP, where rows represent alleles (major or minor allele) and columns represent batches (C5 or C44). Entries of this table reflect allele counts per batch. Subsequently, we performed a Pearson chi-squared test on each table, and thereby obtained a p-value for each SNP. Small p-values indicated that the particular SNP transports batch effects. To correct for multiple testing, we used the Benjamini-Hochberg procedure [73] and filtered all SNPs according to their adjusted p-values, removing SNP’s with an adjusted p-value of less than 0.05. Filtering out 8,664 potentially batch effect signal carrying SNP’s this way, we remained with 4,362,021 SNPs from 22 autosomes.

### Dimensionality reduction

#### GWAS for SNP selection

The dimension of the feature spaces amounts to 4,362,021, agreeing with the number of SNPs that passed quality and batch effect control. On the one hand, this means that the amount of SNPs one can work with is sufficiently large to transport relevant meaning. On the other hand, it means that the number of features is too large for machine learning approaches to not overfit. Therefore, as usual, the dimensionality of the data needs to be reduced for ML approaches to generalize, while preserving the expressiveness of the original set of 4,362,021 features.

To this end, we performed a genome-wide association study using PLINK v1.9 [71], and discarded SNPs that are very unlikely to carry disease status related signals. This reasoning is based on the fact that every SNP must carry an—albeit potentially rather weak—signal in its own right. Using only the training data (see below for descriptions of training, validation and test data), in agreement with the fact that regular GWAS makes use of disease status labels so classifies as part of the training process, we discarded all SNPs whose p-values were greater than 0.05. Note that a threshold of 0.05 is considerably relaxed relative to the stringent threshold of 5 × 10^-8^ [74] that is used in regular GWAS. Unlike in regular GWAS, however, we would like to keep as many *potentially* associated SNPs. So, we do not discard all SNPs whose individual signals are too weak in their own right, knowing that weak individual signals can accumulate to strong signals where SNPs interact in possibly non-additive constellations in the deep learning models that we use. As a result, 505,333 SNPs were retained in this step (see Supplementary Fig. 3); signals of all SNPs discarded were found to be too weak to potentially play a role.

We further annotated all 505,333 SNPs using ANNOVAR (Oct 24, 2019; latest version) [51], assigning them to genes (“gene-based annotation”) based on the human reference genome (hg19). Each SNP could be assigned to least one gene. When annotating SNPs with more than one gene, we kept track of the corresponding mapping relationships: if annotated as “intergenic”, SNPs were assigned to only the nearest gene. If annotated as non-intergenic with variant functions in different genes, we assigned the SNP to all related genes. As a result, SNPs were annotated with 18,279 genes overall, where the vast majority of genes have less than 200 SNPs annotated (see Supplementary Fig. 4).

As usual, we also transformed genotype data according to minor allele information. Each SNP corresponds to a value *i* ∈ {0, 1, 2} in each individual, where *i* is the minor allele count at the particular polymorphic site in the individual.

#### Principal component analysis

Principal component analysis [75, PCA] has been widely applied to exploit SNP data and demonstrated great effectiveness [76]. We recall that whole-genome PCA is not applicable for non-additive approaches to properly work, while gene-based PCA preserves to detect non-additive variant patterns across different genes. In agreement with that reasoning, we performed PCA for each collection of SNPs that became assigned to identical genes. In the following, we refer to that procedure as Gene-PCA.

Correspondingly, for each gene g, we constructed a matrix **A**_*g*_ ∈ {0,1, 2}^*m*×*n*^ where *m* = 10405 is the total number of individual samples, and *n* ∈ 1,…, 1383 corresponds with the number of SNPs that became assigned to gene *g*; note that the maximum amount of SNPs assigned to a gene is 1383, where however the number of genes with more than 200 SNPs assigned was small, see above. See Fig. 1 for an illustration.

Principal component analysis (PCA) can remove noise and generate a robust compressed representation from input features. PCA is efficiently implemented based on singular value decomposition (SVD): **A**_*g*_ is factorized into three matrices

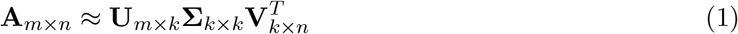

Here **Σ** is a *k* × *k* (usually *k* ≪ *n*) rectangular diagonal matrix of singular values *σ_k_*; importantly, *k* is the number of principal components one will use. **U** and **V** are matrices whose columns are orthogonal unit vectors called the left and the right singular vectors of **A**, respectively. To take into account that the input dimension *n* varies, we varied *k*, relative to *n*, accordingly: *k* = 8 for *n* > 20, *k* = 4 for 4 < *n* ≤ 20 and *k* = 1 for 1 ≤ *n* ≤ 4.

The reduction in dimension we achieve through this procedure is from 505,333 down to 75,584, where each of the 75,584 dimensions corresponds to a principal component computed for one of the 18,279 genes that had become annotated with potentially relevant SNPs, as discussed above.

### The architecture and parameters of DiseaseCapsule

DiseaseCapsule is a neural network model that takes a real-valued vector of length 75,584 as input, and generates binary-valued output, 0 for control/healthy and 1 for ALS/disease. In general, DiseaseCapsule can be flexibly adapted to input vectors of varying sizes, as long as the length of the input vectors does not exceed a certain upper limit; the procedure based on GWAS filtering and Gene-PCA described above warrants this for whole-genome input.

The architecture of DiseaseCapsule follows the architecture of the capsule network that was described in the seminal paper, with modifications to accommodate that the input does not reflect rectangular, pixel-structured image data, and the output to be binary. In detail, DiseaseCapsule consists of three layers: a Fully Connected Layer (FC), a Primary Capsule Layer (PrimaryCaps) and a Phenotype Capsule Layer (PhenoCaps); see Fig. 2 for an illustration and details). The initial fully connected layer replaces the convolutional layers used in the seminal application, reflecting that convolution addresses to process rectangularly arranged image data. However, here we expect interactions between all genes across the whole genome, which the fully connected layer reflects.

The FC layer consists of 150 (regular) neurons followed by ReLU as activation function [77]. During training, a dropout rate of 0.5 was used for FC to prevent overfitting. Correspondingly, the output of FC is a 150 × 1 tensor, so virtually a 150-dimensional vector. This, in turn, is the input for the PrimaryCaps layer.

The PrimaryCaps is the first (“low-level”) capsule layer. As such, it incorporates convolutional operations. It consists of 32 primary capsules. Each of these involves 4 different convolutional filters, implementing a 5 × 1 kernel, operating at a stride of 2, with no zero padding. This means that each convolutional filter computes a 73 × 1 tensor (that is a 73-dimensional vector) from the 150-dimensional input (= FC output) vector. Using 4 convolutional filters per capsule, this results in 73 vectors of length 4 per capsule. This yields 32 × 73 = 2336 vectors of length 4 as output of PrimaryCaps. We refer to each of these vectors as **u**_*i*_, *i* = 1,…, 2336. Further, as is common, one refers to each array of 73 such 4-dimensional vectors as one primary capsule. If needed, we index primary capsules using *k* ∈ {1,…, 32}. Importantly, all 73 **u**_*i*_ making part of one primary capsule *k* share their parameters when transiting to PhenoCaps, the last layer.

Finally, the output of PhenoCaps is used to derive predictions from. PhenoCaps consists of two 16-dimensional vectors **v**_*j*_, *j* = 1, 2, one referring to ‘ALS’ and one to ‘Healthy’. Each phenotype capsule receives input from all 32 primary capsules (i.e., virtually from all 2336**u**_*i*_) making part of PrimaryCaps.

In order to transform PrimaryCaps output into PhenoCaps input, so called pose matrices **W**_*ij*_ are applied to the **w**_*i*_, which yields

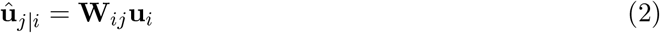

In our case, pose matrices are 4 × 16-dimensional, so the **u**_*j*|*i*_ are 16-dimensional. This corresponds with the dimensionality of PhenoCaps. Pose matrices are learnt during training. As above-mentioned pose matrices are shared for all (73) *i* that refer to an identical primary capsule k. This means that there are 32 pose matrices to be learnt for each *j* = 1, 2.

Subsequently, we performed dynamic routing, as a key feature of capsule networks, intended to improve on the pooling operations. Dynamic routing is an iterative procedure that converges quickly. Here, we used 3 iterations. See Fig. 4 for the corresponding details. According to the routing procedure, the input **s**_*j*_ to one of the PhenoCaps capsules evaluates as

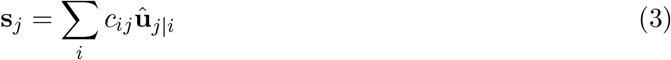

where *c_ij_* are the coupling coefficients, as determined through the dynammic routing procedure. In a rough description, first *b_ij_* are computed by the iterative update 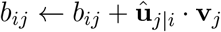 (and initialized as zero), which rewards if **u**_*j*|*i*_ and **v**_*j*_ point in the same direction, which means that the output of primary capsule *i* agrees with phenotype capsule *j*. To turn *b_ij_* into *c_ij_* and ensure that ∑_*i*_ *c_ij_* = 1, one eventually performs

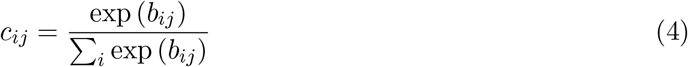

to obtain the coupling coefficients.

Reflecting another key principle of capsule networks, one uses the squashing operation

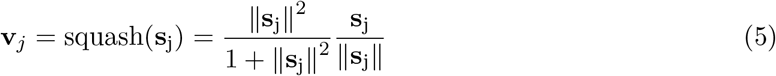

as a non-linear activation function that can process vectors. This ensures that the length of the input vectors **v**_*j*_ is between 0 and 1. Importantly, the length of capsule output vectors **u**_*i*_ and **v**_*j*_ indicates the probability that the capsule “is activated”.

To derive predictions from the **v**_*j*_, categorical cross entropy was employed as the loss function used during training.

### Model training and testing

We randomly split the data set into a set of samples used for training and validation set, that comprised 90% of individuals, and a test set, comprising the remaining 10% of individuals. Importantly, the test set is balanced, that is, the ratio of cases and controls is 1 (here: 520 cases, 520 controls), to ensure that models can be evaluated without misleading biases. See Supplementary Fig. 5 for details.

The entire data set was used for was used for gene based principal component analysis (Gene-PCA). Because dimensionality reduction works in an unsupervised way, so does not require labels, this agrees with a generally applicable protocol. Subsequently, using the training-validation part of the data, five-fold cross validation was performed to optimize the architecture and determine all other hyperparameters of the capsule network (DiseaseCapsule). To ensure a balanced evaluation during validation, validation splits were first randomly selected under the constraint of preserving a ratio of 1:1 between cases and controls. Subsequently, the cases in the remaining training data were upsampled to the same ratio, so as to avoid poor performance in prediction in particular for the minority class (see Supplementary Fig. 5). This reflects a standard procedure in supervised machine learning. Upon having obtained all hyperparameters through cross-validation, the entire training-validation split, upsampled to ensure unbiased training, was used for training. This determines all parameters of the DiseaseCapsule network.

Specifically, we used the Adam algorithm [78] to optimize all parameters in the frame of the usual backpropagation algorithm. We used an initial learning rate of 0.0001, and decayed it by *γ* = 0.8 in each epoch using an exponential scheduler. Optimization ran for 30 epochs, operating at a batch size of 128.

As for the simple three-layer perceptron (MLP) and the basic convolutional neural network (CNN) (consisting of four convolution layers and two dense layers) that we used for benchmarking, see Supplementary Fig. 6 for details. Models were trained for 30 epochs with a batch size of 128 using the Adam optimizer, matching the procedure used for DiseaseCapsule.

### Model interpretation

#### Relating coupling coefficients related with phenotype recognition

When running DiseaseCapsule on the 1040 test samples, coupling coefficients *c_ij_*, *i* = 1,…, 2336, *j* = 1,2 are determined individually for each of the test samples. This is due to the fact that coupling coefficients are determined during the forward step, so are do not correspond to parameters to be learnt during training. According to the general principles of capsule networks, one can interpret coupling coefficients cij as the degree of activation by which **u**_*i*_ contributes to phenotype capsule *j*; large *c_ij_* means that **u**_*i*_ makes a crucial contribution to activating *j*.

As is common, we also determine coupling coefficients *c_kj_* that virtually measure the degree by which primary capsule *k* “activates” phenotype capsule *j*. To obtain *c_kj_* from the *c_ij_*, one sums all coupling coefficients *c_ij_* across all **u**_*i*_ that make part of *k*:

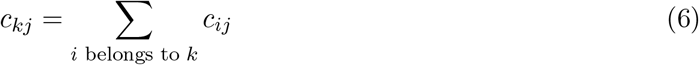

where *j* is either ‘Healthy’ or ‘ALS’, referring to one of the two Phenotype capsules. Recalling that *c_kj_* differ for each individual sample, we eventually average the *c_kj_* across the individuals to obtain a summarizing *c_kj_* one can work with.

#### Selecting and annotating core genes decisive for classification

Evaluating combinations of primary and phenotype capsules according to *c_kj_* from Equation (6) determines primary capsule 5 as the dominant driver to indicate that the phenotype capsule ‘ALS’ gets activated (see Fig. 5a). In other words, genes that significantly contribute to activation of primary capsule 5 are potentially responsible for the development of ALS; in that, these genes are likely to be the predominant factor (see Subsection “Determining genes decisive for classification” in Results). Thanks to capsule networks reflecting non-additive relationships, such genes can interact in arbitrary ways.

Computation of *c_kj_* involves running DiseaseCapsule on all 18,279 genes selected. Here, we would like to determine the genes that play an important role in activating primary capsule 5 in their own right. To do so, we consider each gene *g,g* ∈ (1, 18279) as exclusive input to the trained model. To implement this, we mask all other genes, that is, we set the values of all principal components that do not refer to the particularly picked gene *g* to zero. We do this for each individual sample. Subsequently, we run DiseaseCapsule on all the resulting “one-gene-only” individual samples and note down the resulting coupling coefficients 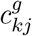, for *k* = 5, *j* = ‘*ALS*’, for each of the individuals. Computation of 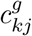 proceeds analogously to Equation (6), when replacing the full individual sample with the “one-gene-only” individual sample. Eventually, also here all individual 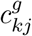 are averaged across the individual samples to obtain a summarizing 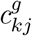 one can work with. We then kept all genes *g* whose 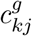 was above the 95-percentile (see Fig. 6a), amounting to 922 genes, as core genes decisive for classification.

In order to annotate the biological functions of these 922 genes, we employed g:Profiler [79] to perform common Gene Ontology (GO) and pathway enrichment analyses.

### Selecting non-additive genes

Let *S* ⊂ *G* be a subset of genes selected from the set *G* of 18,279 genes overall. Let further ACC_DC_(*S*) be the training accuracy achieved by Gene-PCA + DiseaseCapsule (DC) and ACC_LR_(*S*) be the training accuracy of Gene-PCA + LogisticRegression (LR), as the best performing linear approach, when running on only genes *S*. Running DC and LR on S is done by setting values of principal components referring to genes not from *S* to zero, in full analogy as for single genes *g*, as described above.

To determine a good subset of genes that predominantly interact in non-linear constellations, we make use of a genetic algorithm that seeks to determine

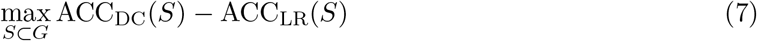

that is the set of genes *S* that delivers the greatest gains in terms of classification performance in the non-linear DC over the the linear LR. To implement the genetic algorithm, we consider subsets of genes as 18279-dimensional binary-valued vectors (*x*_1_, *x*_2_, ⋯, *x*_18279_) ∈ {0,1}^18279^ where *x_g_* = 1 if *g* ∈ *S*, that is gene *g* belongs *t*o S, and *x_g_* =0 otherwise. Representing sets of genes *S* this way, one can implement the common evolutionary operations of genetic algorithms, like ‘selection’, ‘crossover’, ‘mutation’, or ‘fitness evaluation’. For solving the optimization problem Equation (7), we employ ‘Segregative Genetic Algorithms’, as available through the high-performance genetic algorithm toolbox Geatpy v2.6.0 [80].

Subsets *S* were initialized randomly, and the genetic algorithm was run at a population size of 30 and a maximum of generations of 200 as a stopping criterion. The subset of genes S that were determined to maximize Equation (7) can be considered optimal in terms of interacting in exclusively non-additive ways to establish the ‘ALS’ signal in the frame of the DiseaseCapsule network.

### Comparison with other approaches

For a meaningful comparison, we evaluated the performance of various state-of-the-art classification methods. To account for the novel dimensionality reduction protocol ‘Gene-PCA’ we introduced (see above), we ran all methods selected both using ordinary (global) PCA (‘All-PCA’) and ‘Gene-PCA’ for reducing the dimension of the feature space.

As for classification methods themselves, we considered logistic regression and support vector machines (implementing a radial basis function kernel) [81, 82].

We also considered random forests [83] and AdaBoost [84, 85]. Random forests reflect a widely used ensemble learning techniques that is based on collections of decision trees, whose individual predictions are integrated to obtain the ultimate classification. Note that for a random forest to achieve good performance decision trees can be huge, which requires considerable time for training. Here, random forests consist of 100 trees at maximum depth 5, where at most *n* features are considered when looking for the best split with *n* corresponding to the square root of the number of input features. AdaBoost is a boosting algorithm that combines the output of multiple weak learners such as decision trees by assigning more weights to hard-to-classify samples. Even though the final adaptive model is much stronger than the individual classifiers, it can remain sensitive to noisy data and outliers. Here, an AdaBoost model with 1000 decision trees of depth 3 was used. Note that for these conventional machine learning methods, we followed the hyperparameter settings that were determined in our earlier study [46], where identical cohorts were used.

Further, we considered a multilayer perceptron (MLP) and a convolutional neural network (CNN); see Supplementary Fig. 6a and Fig. 6b for the corresponding details. Hyperparameters of models were optimized using cross-validation; performance on the test dataset is reported as final criterion for evaluation.

Evaluating logistic regression without prior dimensionality reduction corresponds to computation of a polygenic risk score (PRS), which is widely used to predict individual trait values from significant genetic markers [86]. while not reducing dimensionality, one can vary the size of the input by application of different significance thresholds *p* for each variant (namely, *p* < 5 × 10^-8^, *p* < 5 × 10^-6^, *p* < 5 × 10^-4^ and *p* < 5 × 10^-2^) where values p are retrieved through standard GWAS.

In addition, we benchmarked several prior methods that are commonly used for raising genomic predictions of complex traits (see review studies [87, 88]), such as Bayesian LASSO (BL) [89], Genomic Best Linear Unbiased Predictor (GBLUP, a ridge-regression type method) [90], Bayesian Ridge Regression (BRR) [91] and BayesB [92]. We performed experiments using the implementations in the R package “BGLR” [93] using default parameters.

Note finally that the application of strategies such as running statistical hypothesis tests on products of variables lack too much power to be applicable in our context, because of the necessary correction for multiple hypothesis testing. The number of pairs of SNP’s amounts to 127.7 billions, while the number of pairs of Gene-PCA’s still amounts to 2.8 billions. We recall as well that hypothesis testing based strategies predominantly cater to the polygenic model, because they are designed to highlight single variants or genes, and so are not able to incorporate more complex functionalities.

### Data preprocessing for Parkinson’s disease

As we did in ALS, we firstly performed an analogous quality control for the data of Parkinson’s disease (PD). Considering the limited number of SNPs in PD data, a lower threshold of minor allele frequency (MAF) was allowed to keep more variants. To be specific, we performed QC using commands “plink –geno 0.1 –maf 0.0001 –hwe 1e-5 midp include-nonctrl” and “plink –het –test-missing midp –pfilter 1e-4” in PD. Only SNPs in autosomal regions were remained. Finally, 135,857 variants and 11402 individuals (5540 cases, 5862 controls) were obtained in PD.

Since the number of remaining SNPs in PD is much fewer than in ALS, there is no need to run the step “GWAS for SNP selection”. Thus, we skipped this step and only performed “Principal component analysis” for dimensionality reduction, which produced the input to various classifiers. As in ALS, we also randomly split the PD data set into a training/validation set (90%) and a test set (10%). Note that the ratio of cases and controls in the test set is balanced.

## Acknowledgements

We would like to thank Bojian Yin and Sumanta Ray for their helpful discussions in the early stage of this project. We thank Marleen Balvert for helpful discussions in the early stages and insightful suggestions when revising the manuscript. We also thank Kristel van der Eijck and Jan Veldink for providing data and helpful information about the mechanisms of ALS. Finally, we thank the International Parkinson’s Disease Genomics Consortium (IPDGC), NINDS, NIA and DoD USAMRAA proposal, 10064005/11348001, for providing the Parkinson’s disease data.

## Funding

XL and XK were supported by the Chinese Scholarship Council. AS was supported by the Dutch ALS Foundation via grant AV20190010. AS was also supported by the Dutch Scientific Organization through Vidi grant 639.072.309 during the early stages of the project. AS further received funding from the European Union’s Horizon 2020 research and innovation programme under Marie Skłodowska-Curie grant agreements No 956229 (ALPACA) and No 872539 (PANGAIA).

## Data availability

The ALS data used in this study has been deposited at dbGaP database (Accession: phs003146.v1.p1). The data of Parkinson’s disease was downloaded from dbGaP Study Accession: phs000918.v1.p1 [52, 53, 54, 55].

## Code availability

The source code of DiseaseCapsule is publicly available on GitHub: https://github.com/HaploKit/ DiseaseCapsule. Other related code for reproducing results and source data for generating figures in this study is publicly available at Zenodo (DOI: 10.5281/zenodo.7118988)[94].

## Competing interests

The authors declare that they have no competing interests.

## Authors’ contributions

XL, XK and AS developed the method. XL and XK implemented the code and conducted the data analysis. XL, XK, and AS wrote the manuscript. All authors read and approved the final version of the manuscript.

## Notes

### Competing Interest Statement

The authors have declared no competing interest.

## References

[1] Giancarlo Logroscino, Bryan J Traynor, Orla Hardiman, Adriano Chiò, Douglas Mitchell, Robert J Swingler, Andrea Millul, Emma Benn, Ettore Beghi, et al. Incidence of amyotrophic lateral sclerosis in europe. Journal of Neurology, Neurosurgery & Psychiatry, 81(4):385–390, 2010.

[2] Kevin Talbot. Motor neuron disease: the bare essentials. Practical neurology, 9(5):303–309, 2009.

[3] Sofie Lautrup, David A Sinclair, Mark P Mattson, and Evandro F Fang. Nad+ in brain aging and neurodegenerative disorders. Cell metabolism, 30(4):630–655, 2019.

[4] JosÉ E de la Rubia, Eraci Drehmer, JosÉ L Platero, MarÍa Benlloch, Jordi Caplliure-Llopis, Carlos Villaron-Casales, Nieves de Bernardo, Jorge AlarcÓn, Cristian Fuente, Sandra Carrera, et al. Efficacy and tolerability of eh301 for amyotrophic lateral sclerosis: a randomized, double-blind, placebo-controlled human pilot study. Amyotrophic Lateral Sclerosis and Frontotemporal Degeneration, 20(1-2):115–122, 2019.

[5] Robert G Miller, Carlayne E Jackson, Edward J Kasarskis, JD England, D Forshew, W Johnston, S Kalra, JS Katz, H Mitsumoto, J Rosenfeld, et al. Practice parameter update: the care of the patient with amyotrophic lateral sclerosis: drug, nutritional, and respiratory therapies (an evidence-based review): report of the quality standards subcommittee of the american academy of neurology. Neurology, 73(15):1218–1226, 2009.

[6] Robert H Brown and Ammar Al-Chalabi. Amyotrophic lateral sclerosis. New England Journal of Medicine, 377(2):162–172, 2017.

[7] Matthew C Kiernan, Steve Vucic, Benjamin C Cheah, Martin R Turner, Andrew Eisen, Orla Hardiman, James R Burrell, and Margaret C Zoing. Amyotrophic lateral sclerosis. The lancet, 377(9769):942–955, 2011.

[8] Ammar Al-Chalabi, Fang Fang, Martha F Hanby, P Nigel Leigh, Christopher E Shaw, Weimin Ye, and Fruhling Rijsdijk. An estimate of amyotrophic lateral sclerosis heritability using twin data. Journal of Neurology, Neurosurgery & Psychiatry, 81(12):1324–1326, 2010.

[9] Philippe A Parone, Sandrine Da Cruz, Joo Seok Han, Melissa McAlonis-Downes, Anne P Vetto, Sandra K Lee, Eva Tseng, and Don W Cleveland. Enhancing mitochondrial calcium buffering capacity reduces aggregation of misfolded sod1 and motor neuron cell death without extending survival in mouse models of inherited amyotrophic lateral sclerosis. Journal of Neuroscience, 33(11):4657–4671, 2013.

[10] Wouter Van Rheenen, Rick AA Van Der Spek, Mark K Bakker, Joke JFA Van Vugt, Paul J Hop, Ramona AJ Zwamborn, Niek De Klein, Harm-Jan Westra, Olivier B Bakker, Patrick Deelen, et al. Common and rare variant association analyses in amyotrophic lateral sclerosis identify 15 risk loci with distinct genetic architectures and neuron-specific biology. Nature genetics, 53(12):1636–1648, 2021.

[11] Hung Phuoc Nguyen, Christine Van Broeckhoven, and Julie van der Zee. Als genes in the genomic era and their implications for ftd. Trends in Genetics, 34(6):404–423, 2018.

[12] Wouter Van Rheenen, Aleksey Shatunov, Annelot M Dekker, Russell L McLaughlin, Frank P Diekstra, Sara L Pulit, Rick AA Van Der Spek, Urmo Võsa, Simone De Jong, Matthew R Robinson, et al. Genome-wide association analyses identify new risk variants and the genetic architecture of amyotrophic lateral sclerosis. Nature genetics, 48(9):1043–1048, 2016.

[13] Marie Ryan, Mark Heverin, Russell L McLaughlin, and Orla Hardiman. Lifetime risk and heritability of amyotrophic lateral sclerosis. JAMA neurology, 76(11):1367–1374, 2019.

[14] Evan A Boyle, Yang I Li, and Jonathan K Pritchard. An expanded view of complex traits: from polygenic to omnigenic. Cell, 169(7):1177–1186, 2017.

[15] Emmanuelle Génin. Missing heritability of complex diseases: case solved? Human Genetics, 139(1):103–113, 2020.

[16] Huwenbo Shi, Gleb Kichaev, and Bogdan Pasaniuc. Contrasting the genetic architecture of 30 complex traits from summary association data. The American Journal of Human Genetics, 99(1):139–153, 2016.

[17] Vivian Tam, Nikunj Patel, Michelle Turcotte, Yohan Bossé, Guillaume Paré, and David Meyre. Benefits and limitations of genome-wide association studies. Nature Reviews Genetics, 20(8):467–484, 2019.

[18] Jason H Moore. The ubiquitous nature of epistasis in determining susceptibility to common human diseases. Human heredity, 56(1-3):73–82, 2003.

[19] Shuo Jiao, Li Hsu, Sonja Berndt, Stéphane Bézieau, Hermann Brenner, Daniel Buchanan, Bette J Caan, Peter T Campbell, Christopher S Carlson, Graham Casey, et al. Genome-wide search for gene-gene interactions in colorectal cancer. PloS one, 7(12):e52535, 2012.

[20] Hung Hung, Yu-Ting Lin, Penweng Chen, Chen-Chien Wang, Su-Yun Huang, and Jung-Ying Tzeng. Detection of gene–gene interactions using multistage sparse and low-rank regression. Biometrics, 72(1):85–94, 2016.

[21] Paola G Ferrario and Inke R König. Transferring entropy to the realm of gxg interactions. Briefings in bioinformatics, 19(1):136–147, 2018.

[22] Sai Zhang, Johnathan Cooper-Knock, Annika K Weimer, Minyi Shi, Tobias Moll, Jack NG Marshall, Calum Harvey, Helia Ghahremani Nezhad, John Franklin, Cleide dos Santos Souza, et al. Genome-wide identification of the genetic basis of amyotrophic lateral sclerosis. Neuron, 2022.

[23] Kurt Hornik, Maxwell Stinchcombe, and Halbert White. Multilayer feedforward networks are universal approximators. Neural networks, 2(5):359–366, 1989.

[24] Guido F Montufar, Razvan Pascanu, Kyunghyun Cho, and Yoshua Bengio. On the number of linear regions of deep neural networks. Advances in neural information processing systems, 27, 2014.

[25] Alex Krizhevsky, Ilya Sutskever, and Geoffrey E Hinton. Imagenet classification with deep convolutional neural networks. Advances in neural information processing systems, 25, 2012.

[26] Yann LeCun, Yoshua Bengio, and Geoffrey Hinton. Deep learning. nature, 521(7553):436–444, 2015.

[27] Laith Alzubaidi, Jinglan Zhang, Amjad J Humaidi, Ayad Al-Dujaili, Ye Duan, Omran Al-Shamma, José Santamaría, Mohammed A Fadhel, Muthana Al-Amidie, and Laith Farhan. Review of deep learning: Concepts, cnn architectures, challenges, applications, future directions. Journal of big Data, 8(1):1–74, 2021.

[28] Karen Simonyan and Andrew Zisserman. Very deep convolutional networks for large-scale image recognition. arXiv preprint arXiv:1409.1556, 2014.

[29] Kaiming He, Xiangyu Zhang, Shaoqing Ren, and Jian Sun. Deep residual learning for image recognition. In Proceedings of the IEEE conference on computer vision and pattern recognition, pages 770–778, 2016.

[30] Gao Huang, Zhuang Liu, Laurens Van Der Maaten, and Kilian Q Weinberger. Densely connected convolutional networks. In Proceedings of the IEEE conference on computer vision and pattern recognition, pages 4700–4708, 2017.

[31] Supriyo Chakraborty, Richard Tomsett, Ramya Raghavendra, Daniel Harborne, Moustafa Alzantot, Federico Cerutti, Mani Srivastava, Alun Preece, Simon Julier, Raghuveer M Rao, et al. Interpretability of deep learning models: A survey of results. In 2017 IEEE smartworld, ubiquitous intelligence & computing, advanced & trusted computed, scalable computing & communications, cloud & big data computing, Internet of people and smart city innovation (smart-world/SCALCOM/UIC/ATC/CBDcom/IOP/SCI), pages 1–6. IEEE, 2017.

[32] Joel Hestness, Sharan Narang, Newsha Ardalani, Gregory Diamos, Heewoo Jun, Hassan Kianinejad, Md Patwary, Mostofa Ali, Yang Yang, and Yanqi Zhou. Deep learning scaling is predictable, empirically. arXiv preprint arXiv:1712.00f09, 2017.

[33] Travers Ching, Daniel S Himmelstein, Brett K Beaulieu-Jones, Alexandr A Kalinin, Brian T Do, Gregory P Way, Enrico Ferrero, Paul-Michael Agapow, Michael Zietz, Michael M Hoffman, et al. Opportunities and obstacles for deep learning in biology and medicine. Journal of The Royal Society Interface, 15(141):20170387, 2018.

[34] Michael Wainberg, Daniele Merico, Andrew Delong, and Brendan J Frey. Deep learning in biomedicine. Nature biotechnology, 36(9):829–838, 2018.

[35] Sara Sabour, Nicholas Frosst, and Geoffrey E Hinton. Dynamic routing between capsules. In Advances in neural information processing systems, pages 3856–3866, 2017.

[36] Sara Sabour, Nicholas Frosst, and Geoffrey Hinton. Matrix capsules with em routing. In 6th international conference on learning representations, ICLR, volume 115, 2018.

[37] Diogo M Camacho, Katherine M Collins, Rani K Powers, James C Costello, and James J Collins. Next-generation machine learning for biological networks. Cell, 173(7):1581–1592, 2018.

[38] Lifei Wang, Rui Nie, Zeyang Yu, Ruyue Xin, Caihong Zheng, Zhang Zhang, Jiang Zhang, and Jun Cai. An interpretable deep-learning architecture of capsule networks for identifying cell-type gene expression programs from single-cell rna-sequencing data. Nature Machine Intelligence, 2(11):693–703, 2020.

[39] Konstantina Kourou, Themis P Exarchos, Konstantinos P Exarchos, Michalis V Karamouzis, and Dimitrios I Fotiadis. Machine learning applications in cancer prognosis and prediction. Computational and structural biotechnology journal, 13:8–17, 2015.

[40] Casimiro A Curbelo Montanez, Paul Fergus, Carl Chalmers, and Jade Hind. Analysis of extremely obese individuals using deep learning stacked autoencoders and genome-wide genetic data. arXiv preprint arXiv:1804.06262, 2018.

[41] Bryan He, Syed Bukhari, Edward Fox, Abubakar Abid, Jeanne Shen, Claudia Kawas, Maria Corrada, Thomas Montine, and James Zou. Ai-enabled in silico immunohistochemical characterization for alzheimer’s disease. Cell reports methods, 2(4):100191, 2022.

[42] Durong Chen, Fuliang Yi, Yao Qin, Jiajia Zhang, Xiaoyan Ge, Hongjuan Han, Jing Cui, Wenlin Bai, Yan Wu, Hongmei Yu, et al. A stacking framework for multi-classification of alzheimer’s disease using neuroimaging and clinical features. Journal of Alzheimer’s Disease, (Preprint):1–10, 2022.

[43] Chenglong Xie, Xu-Xu Zhuang, Zhangming Niu, Ruixue Ai, Sofie Lautrup, Shuangjia Zheng, Yinghui Jiang, Ruiyu Han, Tanima Sen Gupta, Shuqin Cao, et al. Amelioration of alzheimer’s disease pathology by mitophagy inducers identified via machine learning and a cross-species workflow. Nature biomedical engineering, 6(1):76–93, 2022.

[44] Xiong Li, Liyue Liu, Juan Zhou, and Che Wang. Heterogeneity analysis and diagnosis of complex diseases based on deep learning method. Scientific reports, 8(1):1–8, 2018.

[45] Peyton Greenside, Tyler Shimko, Polly Fordyce, and Anshul Kundaje. Discovering epistatic feature interactions from neural network models of regulatory dna sequences. Bioinformatics, 34(17):i629–i637, 2018.

[46] Bojian Yin, Marleen Balvert, Rick AA van der Spek, Bas E Dutilh, Sander Bohte, Jan Veldink, and Alexander Schönhuth. Using the structure of genome data in the design of deep neural networks for predicting amyotrophic lateral sclerosis from genotype. Bioinformatics, 35(14):i538–i547, 2019.

[47] Wayne N Frankel and Nicholas J Schork. Who’s afraid of epistasis? Nature genetics, 14(4):371–373, 1996.

[48] Michael Costanzo, Benjamin VanderSluis, Elizabeth N Koch, Anastasia Baryshnikova, Carles Pons, Guihong Tan, Wen Wang, Matej Usaj, Julia Hanchard, Susan D Lee, et al. A global genetic interaction network maps a wiring diagram of cellular function. Science, 353(6306):aaf1420, 2016.

[49] John L Hartman, Barbara Garvik, and Lee Hartwell. Principles for the buffering of genetic variation. Science, 291(5506):1001–1004, 2001.

[50] Or Zuk, Eliana Hechter, Shamil R Sunyaev, and Eric S Lander. The mystery of missing heritability: Genetic interactions create phantom heritability. Proceedings of the National Academy of Sciences, 109(4):1193–1198, 2012.

[51] Kai Wang, Mingyao Li, and Hakon Hakonarson. Annovar: functional annotation of genetic variants from high-throughput sequencing data. Nucleic acids research, 38(16):e164–e164, 2010.

[52] Paul L Auer, Jill M Johnsen, Andrew D Johnson, Benjamin A Logsdon, Leslie A Lange, Michael A Nalls, Guosheng Zhang, Nora Franceschini, Keolu Fox, Ethan M Lange, et al. Imputation of exome sequence variants into population-based samples and blood-cell-trait-associated loci in african americans: Nhlbi go exome sequencing project. The American Journal of Human Genetics, 91(5):794–808, 2012.

[53] International Parkinson’s Disease Genomics Consortium (IPDGC) and Wellcome Trust Case Control Consortium 2 (WTCCC2). A two-stage meta-analysis identifies several new loci for parkinson’s disease. PLoS genetics, 7(6):e1002142, 2011.

[54] Mike A Nalls, Nathan Pankratz, Christina M Lill, Chuong B Do, Dena G Hernandez, Mohamad Saad, Anita L DeStefano, Eleanna Kara, Jose Bras, Manu Sharma, et al. Large-scale meta-analysis of genome-wide association data identifies six new risk loci for parkinson’s disease. Nature genetics, 46(9):989–993, 2014.

[55] Mike A Nalls, Jose Bras, Dena G Hernandez, Margaux F Keller, Elisa Majounie, Alan E Renton, Mohamad Saad, Iris Jansen, Rita Guerreiro, Steven Lubbe, et al. Neurox, a fast and efficient genotyping platform for investigation of neurodegenerative diseases. Neurobiology of aging, 36(3):1605–e7, 2015.

[56] Mike A Nalls, Cornelis Blauwendraat, Costanza L Vallerga, Karl Heilbron, Sara Bandres-Ciga, Diana Chang, Manuela Tan, Demis A Kia, Alastair J Noyce, Angli Xue, et al. Identification of novel risk loci, causal insights, and heritable risk for parkinson’s disease: a meta-analysis of genome-wide association studies. The Lancet Neurology, 18(12):1091–1102, 2019.

[57] Mariely DeJesus-Hernandez, Ian R Mackenzie, Bradley F Boeve, Adam L Boxer, Matt Baker, Nicola J Rutherford, Alexandra M Nicholson, NiCole A Finch, Heather Flynn, Jennifer Adamson, et al. Expanded ggggcc hexanucleotide repeat in noncoding region of c9orf72 causes chromosome 9p-linked ftd and als. Neuron, 72(2):245–256, 2011.

[58] Elisa Majounie, Alan E Renton, Kin Mok, Elise GP Dopper, Adrian Waite, Sara Rollinson, Adriano Chiò, Gabriella Restagno, Nayia Nicolaou, Javier Simon-Sanchez, et al. Frequency of the c9orf72 hexanucleotide repeat expansion in patients with amy-otrophic lateral sclerosis and frontotemporal dementia: a cross-sectional study. The Lancet Neurology, 11(4):323–330, 2012.

[59] Sónia S Leal and Cláudio M Gomes. Calcium dysregulation links als defective proteins and motor neuron selective vulnerability. Frontiers in cellular neuroscience, 9:225, 2015.

[60] Myrrhe Van Spronsen and Casper C Hoogenraad. Synapse pathology in psychiatric and neurologic disease. Current neurology and neuroscience reports, 10(3):207–214, 2010.

[61] Katarzyna Lepeta, Mychael V Lourenco, Barbara C Schweitzer, Pamela V Martino Adami, Priyanjalee Banerjee, Silvina Catuara-Solarz, Mario de La Fuente Revenga, Alain Marc Guillem, Mouna Haidar, Omamuyovwi M Ijomone, et al. Synaptopathies: synaptic dysfunction in neurological disorders–a review from students to students. Journal of neurochemistry, 138(6):785–805, 2016.

[62] Akito Ikemoto, Shinichi Nakamura, Ichiro Akiguchi, and Asao Hirano. Differential expression between synaptic vesicle proteins and presynaptic plasma membrane proteins in the anterior horn of amyotrophic lateral sclerosis. Acta neuropathologica, 103(2):179–187, 2002.

[63] Katja Burk and R Jeroen Pasterkamp. Disrupted neuronal trafficking in amyotrophic lateral sclerosis. Acta neuropathologica, 137(6):859–877, 2019.

[64] Thomas C Suödhof. Neuroligins and neurexins link synaptic function to cognitive disease. Nature, 455(7215):903–911, 2008.

[65] Antonio Fabregat, Steven Jupe, Lisa Matthews, Konstantinos Sidiropoulos, Marc Gillespie, Phani Garapati, Robin Haw, Bijay Jassal, Florian Korninger, Bruce May, et al. The reactome pathway knowledgebase. Nucleic acids research, 46(D1):D649–D655, 2018.

[66] Kerstin Ure, Hui Lu, Wei Wang, Aya Ito-Ishida, Zhenyu Wu, Ling-jie He, Yehezkel Sztainberg, Wu Chen, Jianrong Tang, and Huda Y Zoghbi. Restoration of mecp2 expression in gabaergic neurons is sufficient to rescue multiple disease features in a mouse model of rett syndrome. Elife, 5:e14198, 2016.

[67] Daniel W Neef, Alex M Jaeger, and Dennis J Thiele. Heat shock transcription factor 1 as a therapeutic target in neurodegenerative diseases. Nature reviews Drug discovery, 10(12):930–944, 2011.

[68] Rebecca San Gil, Lezanne Ooi, Justin J Yerbury, and Heath Ecroyd. The heat shock response in neurons and astroglia and its role in neurodegenerative diseases. Molecular neurodegeneration, 12(1):1–20, 2017.

[69] Alvin Wan, Lisa Dunlap, Daniel Ho, Jihan Yin, Scott Lee, Henry Jin, Suzanne Petryk, Sarah Adel Bargal, and Joseph E Gonzalez. Nbdt: Neural-backed decision trees. arXiv preprint arXiv:2004.00221, 2020.

[70] Project MinE ALS Sequencing Consortium et al. Project mine: study design and pilot analyses of a large-scale whole-genome sequencing study in amyotrophic lateral sclerosis. European Journal of Human Genetics, 26(10):1537, 2018.

[71] Christopher C Chang, Carson C Chow, Laurent CAM Tellier, Shashaank Vattikuti, Shaun M Purcell, and James J Lee. Second-generation plink: rising to the challenge of larger and richer datasets. Gigascience, 4(1):s13742–015, 2015.

[72] Shaun Purcell and Christopher Chang. Plink 1.9. www.cog-genomics.org/plink/1.9/. 2015.

[73] Yoav Benjamini and Yosef Hochberg. Controlling the false discovery rate: a practical and powerful approach to multiple testing. Journal of the Royal statistical society: series B (Methodological), 57(1):289–300, 1995.

[74] International HapMap Consortium et al. A haplotype map of the human genome. Nature, 437(7063):1299, 2005.

[75] Karl Pearson. Liii. on lines and planes of closest fit to systems of points in space. The London, Edinburgh, and Dublin Philosophical Magazine and Journal of Science, 2(11):559–572, 1901.

[76] Alkes L Price, Noah A Zaitlen, David Reich, and Nick Patterson. New approaches to population stratification in genome-wide association studies. Nature Reviews Genetics, 11(7):459–463, 2010.

[77] Vinod Nair and Geoffrey E Hinton. Rectified linear units improve restricted boltzmann machines. In Icml, 2010.

[78] Diederik P Kingma and Jimmy Ba. Adam: A method for stochastic optimization. arXiv preprint arXiv:1412.6980, 2014.

[79] Uku Raudvere, Liis Kolberg, Ivan Kuzmin, Tambet Arak, Priit Adler, Hedi Peterson, and Jaak Vilo. g: Profiler: a web server for functional enrichment analysis and conversions of gene lists (2019 update). Nucleic acids research, 47(W1):W191–W198, 2019.

[80] et.al. Jazzbin. geatpy: The genetic and evolutionary algorithm toolbox with high performance in python, 2020.

[81] Thorsten Joachims. Text categorization with support vector machines: Learning with many relevant features. In European conference on machine learning, pages 137–142. Springer, 1998.

[82] Vladimir Vapnik. The support vector method of function estimation. In Nonlinear Modeling, pages 55–85. Springer, 1998.

[83] Leo Breiman. Random forests. Machine learning, 45(1):5–32, 2001.

[84] Yoav Freund, Robert Schapire, and Naoki Abe. A short introduction to boosting. Journal-Japanese Society For Artificial Intelligence, 14(771-780):1612, 1999.

[85] Jerome Friedman, Trevor Hastie, Robert Tibshirani, et al. Additive logistic regression: a statistical view of boosting (with discussion and a rejoinder by the authors). The annals of statistics, 28(2):337–407, 2000.

[86] Frank Dudbridge. Power and predictive accuracy of polygenic risk scores. PLoS genetics, 9(3), 2013.

[87] Pau Bellot, Gustavo de Los Campos, and Miguel Pérez-Enciso. Can deep learning improve genomic prediction of complex human traits? Genetics, 210(3):809–819, 2018.

[88] M Pérez-Enciso and LM Zingaretti. A guide for using deep learning for complex trait genomic prediction. genes (basel) 10: 553, 2019.

[89] Trevor Park and George Casella. The bayesian lasso. Journal of the American Statistical Association, 103(482):681–686, 2008.

[90] Gustavo de Los Campos, Ana I Vazquez, Rohan Fernando, Yann C Klimentidis, and Daniel Sorensen. Prediction of complex human traits using the genomic best linear unbiased predictor. PLoS genetics, 9(7):e1003608, 2013.

[91] CR Henderson. Applications of linear models in animal breeding (university of guelph, guelph, on, canada). Applications of linear models in animal breeding. University of Guelph, Guelph, ON, Canada, 1984.

[92] Theo HE Meuwissen, Ben J Hayes, and ME1461589 Goddard. Prediction of total genetic value using genome-wide dense marker maps. genetics, 157(4):1819–1829, 2001.

[93] Paulino Pérez and Gustavo de Los Campos. Genome-wide regression and prediction with the bglr statistical package. Genetics, 198(2):483–495, 2014.

[94] Luo, X. and Kang, X. and Schönhuth, A. Diseasecapsule: v1.0.0. zenodo, 2022.

