## Supplementary Material for "Predicting the prevalence of complex genetic diseases from individual genotype profiles using capsule networks"

### Supplementary Information

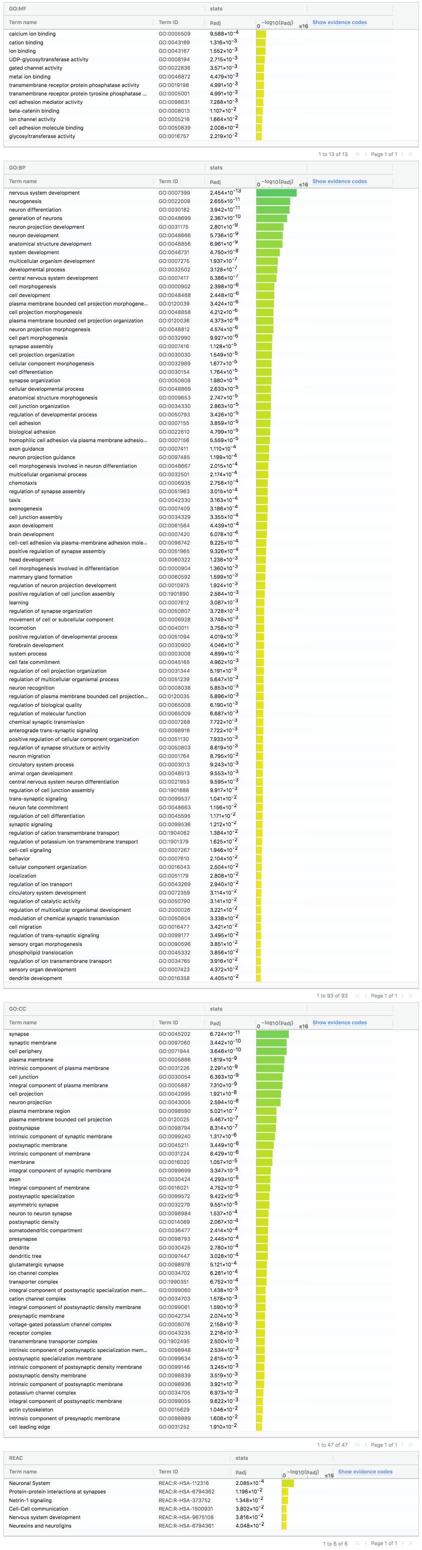

GO:MF

| Term name | Term ID | Padj |
|---|---|---|
| calcium ion binding | GO:0005509 | $9.588\times10^{-4}$ |
| cation binding | GO:0043169 | $1.316\times10^{-3}$ |
| ion binding | GO:0043167 | $1.552\times10^{-3}$ |
| UDP-glycosyltransferase activity | GO:0008194 | $2.715\times10^{-3}$ |
| gated channel activity | GO:0022836 | $3.571\times10^{-3}$ |
| metal ion binding | GO:0046872 | $4.479\times10^{-3}$ |
| transmembrane receptor protein phosphatase activity | GO:0019198 | $4.991\times10^{-3}$ |
| transmembrane receptor protein tyrosine phosphatase ... | GO:0005001 | $4.991\times10^{-3}$ |
| cell adhesion mediator activity | GO:0098631 | $7.288\times10^{-3}$ |
| beta-catenin binding | GO:0008013 | $1.107\times10^{-2}$ |
| ion channel activity | GO:0005216 | $1.664\times10^{-2}$ |
| cell adhesion molecule binding | GO:0050839 | $2.008\times10^{-2}$ |
| glycosyltransferase activity | GO:0016757 | $2.219\times10^{-2}$ |

1 to 13 of 13 Page 1 of 1

GO:BP

| Term name | Term ID | Padj |
|---|---|---|
| nervous system development | GO:0007399 | $2.454\times10^{-13}$ |
| neurogenesis | GO:0022008 | $2.655\times10^{-11}$ |
| neuron differentiation | GO:0030182 | $3.942\times10^{-11}$ |
| generation of neurons | GO:0048699 | $2.367\times10^{-10}$ |
| neuron projection development | GO:0031175 | $2.801\times10^{-9}$ |
| neuron development | GO:0048666 | $5.736\times10^{-9}$ |
| anatomical structure development | GO:0048856 | $6.961\times10^{-9}$ |
| system development | GO:0048731 | $4.750\times10^{-8}$ |
| multicellular organism development | GO:0007275 | $1.937\times10^{-7}$ |
| developmental process | GO:0032502 | $3.128\times10^{-7}$ |
| central nervous system development | GO:0007417 | $5.386\times10^{-7}$ |
| cell morphogenesis | GO:0000902 | $2.398\times10^{-6}$ |
| cell development | GO:0048468 | $2.448\times10^{-6}$ |
| plasma membrane bounded cell projection morphogene... | GO:0120039 | $3.424\times10^{-6}$ |
| cell projection morphogenesis | GO:0048858 | $4.212\times10^{-6}$ |
| plasma membrane bounded cell projection organization | GO:0120036 | $4.373\times10^{-6}$ |
| neuron projection morphogenesis | GO:0048812 | $4.574\times10^{-6}$ |
| cell part morphogenesis | GO:0032990 | $9.927\times10^{-6}$ |
| synapse assembly | GO:0007416 | $1.128\times10^{-5}$ |
| cell projection organization | GO:0030030 | $1.549\times10^{-5}$ |
| cellular component morphogenesis | GO:0032989 | $1.677\times10^{-5}$ |
| cell differentiation | GO:0030154 | $1.764\times10^{-5}$ |
| synapse organization | GO:0050808 | $1.980\times10^{-5}$ |
| cellular developmental process | GO:0048869 | $2.633\times10^{-5}$ |
| anatomical structure morphogenesis | GO:0009653 | $2.747\times10^{-5}$ |
| cell junction organization | GO:0034330 | $2.863\times10^{-5}$ |
| regulation of developmental process | GO:0050793 | $3.426\times10^{-5}$ |
| cell adhesion | GO:0007155 | $3.859\times10^{-5}$ |
| biological adhesion | GO:0022610 | $4.799\times10^{-5}$ |
| homophilic cell adhesion via plasma membrane adhesio... | GO:0007156 | $5.559\times10^{-5}$ |
| axon guidance | GO:0007411 | $1.110\times10^{-4}$ |
| neuron projection guidance | GO:0097485 | $1.199\times10^{-4}$ |
| cell morphogenesis involved in neuron differentiation | GO:0048667 | $2.015\times10^{-4}$ |
| multicellular organismal process | GO:0032501 | $2.174\times10^{-4}$ |
| chemotaxis | GO:0006935 | $2.758\times10^{-4}$ |
| regulation of synapse assembly | GO:0051963 | $3.015\times10^{-4}$ |
| taxis | GO:0042330 | $3.163\times10^{-4}$ |
| axonogenesis | GO:0007409 | $3.186\times10^{-4}$ |
| cell junction assembly | GO:0034329 | $3.355\times10^{-4}$ |
| axon development | GO:0061564 | $4.439\times10^{-4}$ |
| brain development | GO:0007420 | $5.078\times10^{-4}$ |
| cell-cell adhesion via plasma-membrane adhesion mole... | GO:0098742 | $8.225\times10^{-4}$ |
| positive regulation of synapse assembly | GO:0051965 | $9.326\times10^{-4}$ |
| head development | GO:0060322 | $1.238\times10^{-3}$ |
| cell morphogenesis involved in differentiation | GO:0000904 | $1.360\times10^{-3}$ |
| mammary gland formation | GO:0060592 | $1.599\times10^{-3}$ |
| regulation of neuron projection development | GO:0010975 | $1.924\times10^{-3}$ |
| positive regulation of cell junction assembly | GO:1901890 | $2.584\times10^{-3}$ |
| learning | GO:0007612 | $3.087\times10^{-3}$ |
| regulation of synapse organization | GO:0050807 | $3.728\times10^{-3}$ |
| movement of cell or subcellular component | GO:0006928 | $3.749\times10^{-3}$ |
| locomotion | GO:0040011 | $3.758\times10^{-3}$ |
| positive regulation of developmental process | GO:0051094 | $4.019\times10^{-3}$ |
| forebrain development | GO:0030900 | $4.046\times10^{-3}$ |
| system process | GO:0003008 | $4.899\times10^{-3}$ |
| cell fate commitment | GO:0045165 | $4.962\times10^{-3}$ |
| regulation of cell projection organization | GO:0031344 | $5.191\times10^{-3}$ |
| regulation of multicellular organismal process | GO:0051239 | $5.647\times10^{-3}$ |
| neuron recognition | GO:0008038 | $5.853\times10^{-3}$ |
| regulation of plasma membrane bounded cell projection... | GO:0120035 | $5.896\times10^{-3}$ |
| regulation of biological quality | GO:0065008 | $6.190\times10^{-3}$ |
| regulation of molecular function | GO:0065009 | $6.687\times10^{-3}$ |
| chemical synaptic transmission | GO:0007268 | $7.722\times10^{-3}$ |
| anterograde trans-synaptic signaling | GO:0098916 | $7.722\times10^{-3}$ |
| positive regulation of cellular component organization | GO:0051130 | $7.933\times10^{-3}$ |
| regulation of synapse structure or activity | GO:0050803 | $8.619\times10^{-3}$ |
| neuron migration | GO:0001764 | $8.795\times10^{-3}$ |
| circulatory system process | GO:0003013 | $9.243\times10^{-3}$ |
| animal organ development | GO:0048513 | $9.553\times10^{-3}$ |
| central nervous system neuron differentiation | GO:0021953 | $9.595\times10^{-3}$ |
| regulation of cell junction assembly | GO:1901888 | $9.917\times10^{-3}$ |
| trans-synaptic signaling | GO:0099537 | $1.041\times10^{-2}$ |
| neuron fate commitment | GO:0048663 | $1.156\times10^{-2}$ |
| regulation of cell differentiation | GO:0045595 | $1.171\times10^{-2}$ |
| synaptic signaling | GO:0099536 | $1.212\times10^{-2}$ |
| regulation of cation transmembrane transport | GO:1904062 | $1.384\times10^{-2}$ |
| regulation of potassium ion transmembrane transport | GO:1901379 | $1.625\times10^{-2}$ |
| cell-cell signaling | GO:0007267 | $1.946\times10^{-2}$ |
| behavior | GO:0007610 | $2.104\times10^{-2}$ |
| cellular component organization | GO:0016043 | $2.504\times10^{-2}$ |
| localization | GO:0051179 | $2.808\times10^{-2}$ |
| regulation of ion transport | GO:0043269 | $2.940\times10^{-2}$ |
| circulatory system development | GO:0072359 | $3.114\times10^{-2}$ |
| regulation of catalytic activity | GO:0050790 | $3.141\times10^{-2}$ |
| regulation of multicellular organismal development | GO:2000026 | $3.221\times10^{-2}$ |
| modulation of chemical synaptic transmission | GO:0050804 | $3.338\times10^{-2}$ |
| cell migration | GO:0016477 | $3.421\times10^{-2}$ |
| regulation of trans-synaptic signaling | GO:0099177 | $3.495\times10^{-2}$ |
| sensory organ morphogenesis | GO:0090596 | $3.851\times10^{-2}$ |
| phospholipid translocation | GO:0045332 | $3.856\times10^{-2}$ |
| regulation of ion transmembrane transport | GO:0034765 | $3.916\times10^{-2}$ |
| sensory organ development | GO:0007423 | $4.372\times10^{-2}$ |
| dendrite development | GO:0016358 | $4.405\times10^{-2}$ |

1 to 93 of 93 Page 1 of 1

GO:CC

| Term name | Term ID | Padj |
|---|---|---|
| synapse | GO:0045202 | $6.724\times10^{-11}$ |
| synaptic membrane | GO:0097060 | $3.442\times10^{-10}$ |
| cell periphery | GO:0071944 | $3.646\times10^{-10}$ |
| plasma membrane | GO:0005886 | $1.819\times10^{-9}$ |
| intrinsic component of plasma membrane | GO:0031226 | $2.291\times10^{-9}$ |
| cell junction | GO:0030054 | $6.393\times10^{-9}$ |
| integral component of plasma membrane | GO:0005887 | $7.310\times10^{-9}$ |
| cell projection | GO:0042995 | $1.921\times10^{-8}$ |
| neuron projection | GO:0043005 | $2.594\times10^{-8}$ |
| plasma membrane region | GO:0098590 | $5.021\times10^{-7}$ |
| plasma membrane bounded cell projection | GO:0120025 | $5.467\times10^{-7}$ |
| postsynapse | GO:0098794 | $8.314\times10^{-7}$ |
| intrinsic component of synaptic membrane | GO:0099240 | $1.317\times10^{-6}$ |
| postsynaptic membrane | GO:0045211 | $3.449\times10^{-6}$ |
| intrinsic component of membrane | GO:0031224 | $8.429\times10^{-6}$ |
| membrane | GO:0016020 | $1.057\times10^{-5}$ |
| integral component of synaptic membrane | GO:0099699 | $3.347\times10^{-5}$ |
| axon | GO:0030424 | $4.293\times10^{-5}$ |
| integral component of membrane | GO:0016021 | $4.752\times10^{-5}$ |
| postsynaptic specialization | GO:0099572 | $9.422\times10^{-5}$ |
| asymmetric synapse | GO:0032279 | $9.551\times10^{-5}$ |
| neuron to neuron synapse | GO:0098984 | $1.537\times10^{-4}$ |
| postsynaptic density | GO:0014069 | $2.067\times10^{-4}$ |
| somatodendritic compartment | GO:0036477 | $2.414\times10^{-4}$ |
| presynapse | GO:0098793 | $2.445\times10^{-4}$ |
| dendrite | GO:0030425 | $2.780\times10^{-4}$ |
| dendritic tree | GO:0097447 | $3.028\times10^{-4}$ |
| glutamatergic synapse | GO:0098978 | $5.121\times10^{-4}$ |
| ion channel complex | GO:0034702 | $6.281\times10^{-4}$ |
| transporter complex | GO:1990351 | $6.752\times10^{-4}$ |
| integral component of postsynaptic specialization mem... | GO:0099060 | $1.438\times10^{-3}$ |
| cation channel complex | GO:0034703 | $1.578\times10^{-3}$ |
| integral component of postsynaptic density membrane | GO:0099061 | $1.590\times10^{-3}$ |
| presynaptic membrane | GO:0042734 | $2.074\times10^{-3}$ |
| voltage-gated potassium channel complex | GO:0008076 | $2.158\times10^{-3}$ |
| receptor complex | GO:0043235 | $2.216\times10^{-3}$ |
| transmembrane transporter complex | GO:1902495 | $2.500\times10^{-3}$ |
| intrinsic component of postsynaptic specialization mem... | GO:0098948 | $2.534\times10^{-3}$ |
| postsynaptic specialization membrane | GO:0099634 | $2.615\times10^{-3}$ |
| intrinsic component of postsynaptic density membrane | GO:0099146 | $3.245\times10^{-3}$ |
| postsynaptic density membrane | GO:0098839 | $3.519\times10^{-3}$ |
| intrinsic component of postsynaptic membrane | GO:0098936 | $3.921\times10^{-3}$ |
| potassium channel complex | GO:0034705 | $6.973\times10^{-3}$ |
| integral component of postsynaptic membrane | GO:0099055 | $9.622\times10^{-3}$ |
| actin cytoskeleton | GO:0015629 | $1.046\times10^{-2}$ |
| intrinsic component of presynaptic membrane | GO:0098889 | $1.606\times10^{-2}$ |
| cell leading edge | GO:0031252 | $1.910\times10^{-2}$ |

1 to 47 of 47 Page 1 of 1

REAC

| Term name | Term ID | Padj |
|---|---|---|
| Neuronal System | REAC:R-HSA-112316 | $2.085\times10^{-4}$ |
| Protein-protein interactions at synapses | REAC:R-HSA-6794362 | $1.196\times10^{-2}$ |
| Netrin-1 signaling | REAC:R-HSA-373752 | $1.348\times10^{-2}$ |
| Cell-Cell communication | REAC:R-HSA-1500931 | $3.802\times10^{-2}$ |
| Nervous system development | REAC:R-HSA-9675108 | $3.816\times10^{-2}$ |
| Neurexins and neuroligins | REAC:R-HSA-6794361 | $4.048\times10^{-2}$ |

1 to 6 of 6 Page 1 of 1

**Supplementary Figure 1.** Functional annotation of 922 genes decisive for classification.

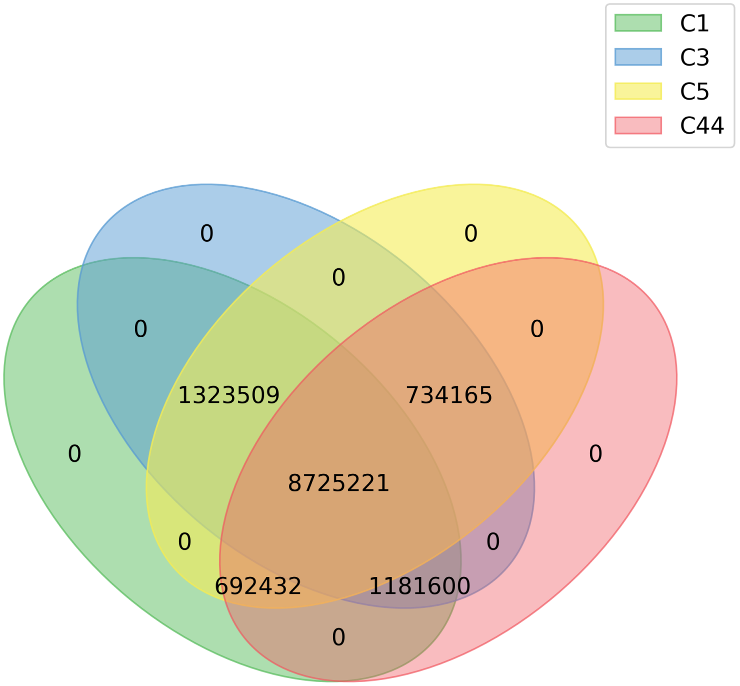

**Supplementary Figure 2.** Venn plot for not quality controlled SNPs called per batch.

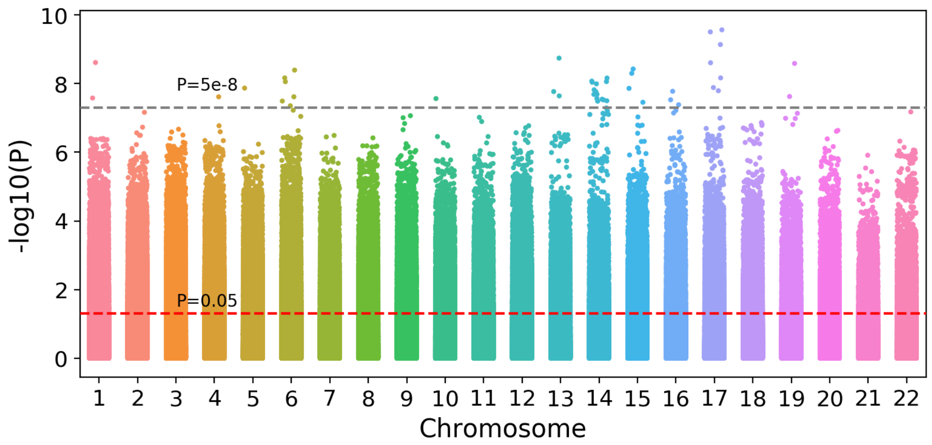

**Supplementary Figure 3.** Manhattan plot for GWAS SNPs. The grey dashed line represents the standard GWAS threshold, and red dashed line represents a more relaxed threshold (0.05).

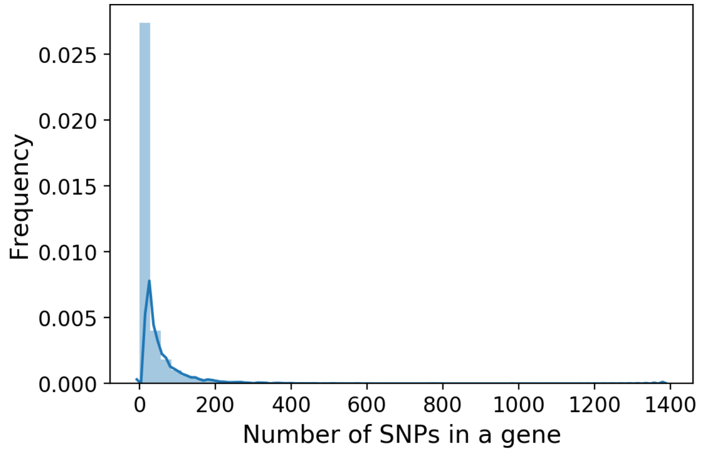

**Supplementary Figure 4.** Distribution for the number of SNPs in genes.

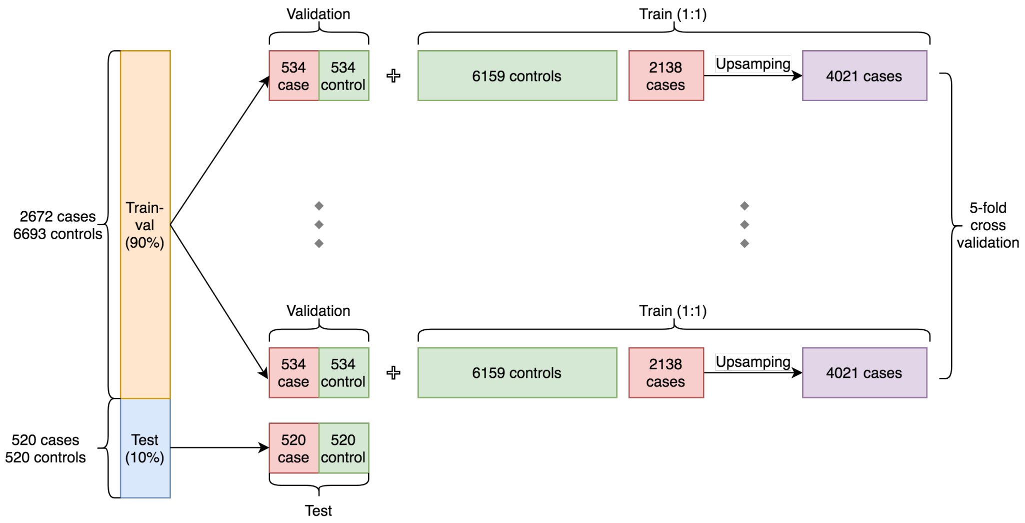

**Supplementary Figure 5.** Split dataset for model training and testing.

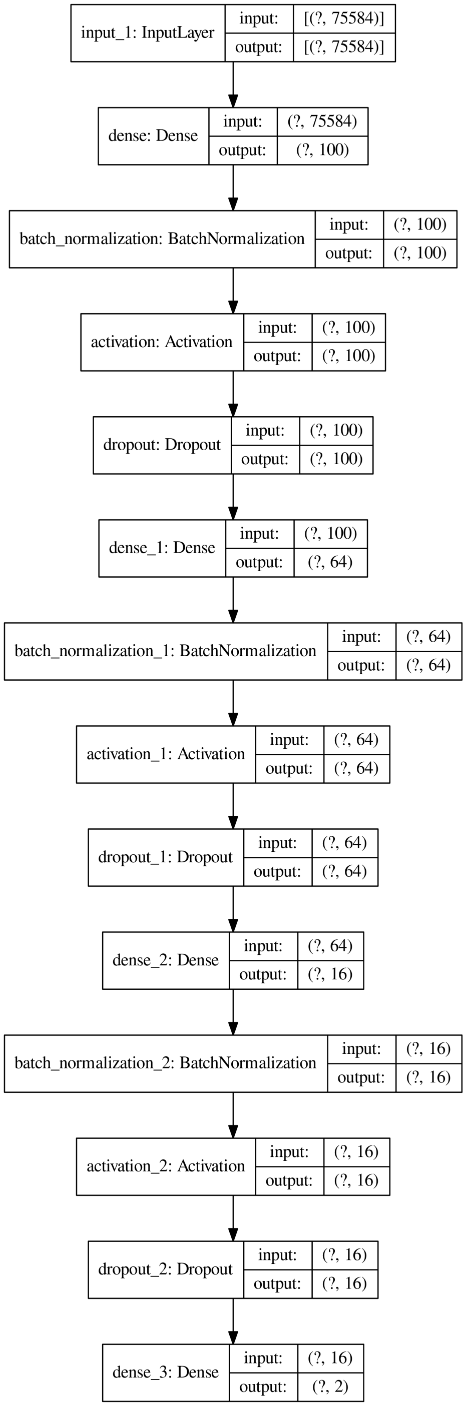

**(a)** The model graph of MLP

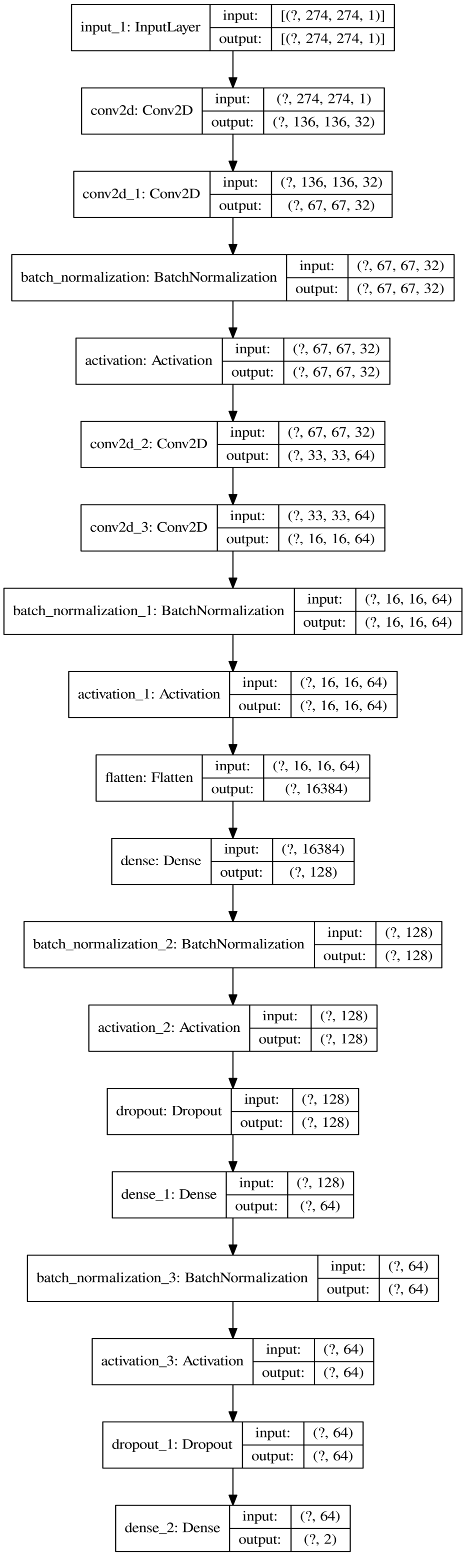

**(b)** The model graph of CNN

**Supplementary Figure 6.** The visualization of other DNN models.

| Dimension reduction | Classifier | Accuracy | Precision | Recall | F1-Score |
|---|---|---|---|---|---|
| Gene-AE | LogisticRegression | $83.4 \pm 0.1$ | $92.6 \pm 0.3$ | $72.5 \pm 0.4$ | $81.3 \pm 0.2$ |
| Gene-AE | MLP | $83.2 \pm 0.5$ | $92.4 \pm 0.5$ | $72.4 \pm 1.0$ | $81.2 \pm 0.7$ |
| Gene-AE | SVM | $82.1 \pm 0.0$ | $93.9 \pm 0.0$ | $68.7 \pm 0.0$ | $79.3 \pm 0.0$ |
| Gene-AE | DiseaseCapsule | $79.6 \pm 7.1$ | $91.5 \pm 3.5$ | $66.1 \pm 18.0$ | $75.0 \pm 12.9$ |
| Gene-AE | CNN | $74.0 \pm 1.0$ | $85.4 \pm 1.0$ | $57.9 \pm 1.8$ | $69.0 \pm 1.5$ |
| Gene-AE | RandomForest | $63.4 \pm 0.9$ | $72.1 \pm 1.7$ | $43.7 \pm 1.6$ | $54.4 \pm 1.4$ |

**Supplementary Table 1.** Classification results for test dataset using autoencoders for dimension reduction. Values are represented as mean ± sd (%), which are calculated by repeating training-testing process for ten times with the same train/test dataset and hyperparameters. Gene-AE: using autoencoder for dimension reduction on the SNP profile of each individual gene. The number of neurons in the autoencoder is 128, 64, 32 from peripheral layer to the middle layer in both encoder and decoder, the number of neurons in the bottleneck layer is 8 or 4 or 1, which depends on the length of the input, i.e. the number of SNPs (processing with the same way in Gene-PCA).

| Dimension reduction | Classifier | Accuracy | Precision | Recall | F1-Score |
|---|---|---|---|---|---|
| Gene-PCA | BL | 69.0 | 91.3 | 38.3 | 54.0 |
| Gene-PCA | GBLUP | 68.9 | 91.3 | 38.1 | 53.8 |
| Gene-PCA | BRR | 68.8 | 91.6 | 37.7 | 53.4 |
| Gene-PCA | BayesB | 67.0 | 91.7 | 33.5 | 49.1 |
| All-PCA | BL | 69.7 | 87.4 | 42.2 | 56.9 |
| All-PCA | BayesB | 69.6 | 86.4 | 42.6 | 57.1 |
| All-PCA | BRR | 68.4 | 91.5 | 36.9 | 52.6 |
| All-PCA | GBLUP | 67.9 | 91.2 | 35.7 | 51.3 |
| - | $BRR^a$ | 66.4 | 93.9 | 31.2 | 46.8 |
| - | $BRR^b$ | 62.4 | 98.1 | 21.1 | 34.7 |

**Supplementary Table 2.** Classification results of alternative methods on the ALS test dataset. The values are represented as percentages. BL: Bayesian LASSO, GBLUP: Genomic Best Linear Unbiased Predictor, BRR: Bayesian Ridge Regression. $a$,$b$ used the original SNPs(without dimension reduction) that were selected by GWAS with the threshold ($P < 5 \times 10^{-4}$, $P < 5 \times 10^{-2}$) respectively.

| Dimension reduction | Classifier | Accuracy | Precision | Recall | F1-Score |
|---|---|---|---|---|---|
| Gene-PCA | DiseaseCapsule | **62.0** | 60.7 | **68.1** | **64.2** |
| Gene-PCA | MLP | 60.6 | 60.7 | 60.4 | 60.5 |
| Gene-PCA | SVM | 59.6 | 60.5 | 55.4 | 57.8 |
| Gene-PCA | RandomForest | 58.6 | 57.9 | 62.8 | 60.3 |
| Gene-PCA | LogisticRegression | 58.4 | 58.7 | 56.8 | 57.7 |
| Gene-PCA | CNN | 56.0 | 56.3 | 53.3 | 54.8 |
| All-PCA | SVM | 61.4 | **62.2** | 58.2 | 60.1 |
| All-PCA | LogisticRegression | 59.5 | 59.8 | 57.9 | 58.8 |
| All-PCA | DiseaseCapsule | 57.5 | 59.0 | 49.5 | 53.8 |
| All-PCA | MLP | 57.2 | 57.9 | 52.6 | 55.1 |
| All-PCA | RandomForest | 51.1 | 55.3 | 11.1 | 18.5 |
| All-PCA | CNN | 50.2 | 50.2 | 43.0 | 46.4 |
| - | Polygenic Risk Score[a] | 59.9 | 60.6 | 56.8 | 58.6 |

**Supplementary Table 3.** Classification results for Parkinson's disease test data. The values are represented as percentages. MLP: multilayer perceptron, SVM: support vector machine, CNN: convolutional neural network. $a$ represent the PRS-based model that the SNPs were selected by GWAS with the threshold ($P < 5 \times 10^{-2}$).

| | | | | | | | | | |
|---|---|---|---|---|---|---|---|---|---|
| NALCN-AS1 | PCAT29 | SPACA1 | ATXN8OS | CADPS | CNBD1 | CST1 | CST2 | DCC | DEFB112 |
| LINC00917 | LINC01741 | LINC02433 | LINC02444 | **LIPC** | LOC101927915 | LOC102724265 | LRRC3B | MAST4 | MIR4268 |
| MIR4432HG | MROH5 | P3H2 | PLCG2 | ROBO2 | SMARCA2 | TAFA2 | ACKR3 | ADGRL2 | AGBL4 |
| ANKRD33B | ANO5 | ANXA1 | APOL6 | ASAP2 | ATP9A | AUTS2 | CACNA1E | CALCA | CAND1 |
| CD36 | CDH18 | CDH7 | CHSY3 | **CNTN4** | **CNTN6** | CSMD1 | CT66 | DLEU1 | DLGAP2 |
| DOCK2 | ELMO1 | EYA1 | FMN1 | GABBR2 | GCNT2 | HAS2 | HDAC4 | HLA-DPB2 | HS3ST4 |
| HS6ST3 | IPO5 | KCNE4 | KCNK1 | KCTD8 | KDM4C | LINC00276 | LINC00358 | LINC00593 | LINC00607 |
| LINC01243 | LINC01248 | LINC01362 | LINC01570 | LINC01619 | LINC01980 | LINC02093 | LINC02261 | LINC02364 | LINC02374 |
| LINC02465 | LINC02468 | LINC02500 | LINC02615 | LOC101927987 | LOC105370980 | MACROD2 | MAGI1 | MAP3K7 | MED13L |
| MIR7702 | MMUT | MYO10 | NAALADL2 | NFATC2 | NOS1AP | NRG3 | NUP35 | NXPH1 | OR52H1 |
| OSR1 | PCAT5 | PER4 | PPP2R2C | PPP2R3A | PTCHD4 | PTPRE | PTPRN2 | RAPH1 | RASSF3 |
| RBMS3 | RHBDD1 | RNLS | ROR1 | RORA | SASH1 | SETD3 | SLC8A1 | SMIM31 | SNTN |
| SNX2 | SPAG17 | SPATA31C2 | STARD13 | STIM2 | SUFU | TAB2 | TBC1D30 | TECTB | TFAP2A |
| TFRC | THNSL2 | TLE1 | TLE4 | TMEM132D | TRIM33 | TRPC6 | TRPS1 | TSNARE1 | TUSC3 |
| TYRP1 | UGGT2 | UNC93A | ZHX2 | ZNF283 | ZNF518B | AACSP1 | ABCC13 | ACTL7B | ADAMTS17 |
| ADAMTS5 | ADCY8 | ADGB | ADGRB3 | AJAP1 | AKAP6 | ARHGAP28 | ARL4A | ASAP1 | ASIC2 |
| ATP10D | C11orf74 | C3orf56 | C8orf34 | CA6 | CA8 | CACNB4 | CAMKMT | CBLB | CCR1 |
| CCR2 | CD2AP | CDH8 | CDKAL1 | CDYL2 | CLSTN2 | CMC1 | CMYA5 | CNTN1 | COL4A4 |
| COL6A3 | CPE | CRYL1 | CT69 | CTAGE1 | CYLC2 | DLG2 | DMAC1 | DOCK1 | DOCK4 |
| **DPP6** | DYRK1A | EEF2KMT | EMILIN2 | **ERBB4** | ETV1 | EXOC6B | F11-AS1 | FAM241A | FHIT |
| FOXP1 | FSHR | FSIP2-AS2 | FSTL5 | FYCO1 | GALNT17 | GALNTL6 | GLT1D1 | GNS | GPC5 |
| GPR12 | GRIA1 | GSTA2 | HS3ST3A1 | IMMP2L | INSR | IPMK | JAG2 | KBTBD11 | KCNAB1 |
| KCNK18 | KCNMA1 | KIF4B | LARGE1 | LBH | LCORL | LHPP | LINC00410 | LINC00616 | LINC00838 |
| LINC00924 | LINC00963 | LINC00970 | LINC01010 | LINC01148 | LINC01170 | LINC01239 | LINC01256 | LINC01257 | LINC01324 |
| LINC01411 | LINC01445 | LINC01559 | LINC01571 | LINC01622 | LINC01648 | LINC01680 | LINC01682 | LINC01720 | LINC01799 |
| LINC01826 | LINC01841 | LINC01941 | LINC01947 | LINC01951 | LINC02008 | LINC02054 | LINC02103 | LINC02201 | LINC02208 |
| LINC02254 | LINC02393 | LINC02474 | LINC02485 | LINC02549 | LINC02645 | LINC02667 | LINGO2 | LMNB1-DT | LOC101927066 |
| LOC101929297 | LOC101929563 | LOC102723427 | LOC102723883 | LOC105376360 | LOC646736 | LRP1B | LRRN1 | LYZL1 | MAGI2 |
| MAP3K21 | MEAT6 | MED15 | MEIS2 | MICAL3 | MINDY3 | MIR100HG | MIR1-1HG-AS1 | MIR12113 | MIR1263 |
| MIR3974 | MIR4318 | MIR4422HG | MIR548AB | MIR548AP | MIR548XHG | MIR5702 | MOB3B | MROH7-TTC4 | MROH9 |
| NAV3 | NCAM2 | NEBL | NEDD4L | NEK1 | NELL1 | NPR3 | NRIP1 | NRXN1 | NRXN3 |
| NSMCE2 | NTPCR | OAT | OLIG3 | OR52M1 | PACRG | PACS1 | PALLD | PALM2-AKAP2 | PDGFD |
| PGM2 | PKIA | PLPPR1 | POMGNT2 | PRUNE2 | PSD3 | **RAMP3** | ROBO1 | RPSA | RTN1 |
| RXFP2 | SALL3 | SATB2 | SEC61A2 | SEZ6L | SGCD | SLC2A9 | SLC45A4 | SLC5A4-AS1 | SLCO3A1 |
| SLITRK6 | SNTB1 | SNX24 | SNX29 | SORCS2 | SPATA16 | SPIDR | SPOCK1 | SPRED2 | STX8 |
| STXBP6 | SUMF1 | SYN3 | SYT4 | TAFA1 | TAFA5 | TEMN3-AS1 | THEG5 | THEM4 | TIFA |
| TLR4 | TLR5 | TMCC3 | TMEM135 | TMEM178B | TOX | TP63 | TRAM1L1 | TRAPPC9 | TRPC4 |
| TUT4 | UNC13C | USP12 | VPS50 | WASL | WDR36 | XXYLT1 | YIPF7 | ZBTB7C | ZFP42 |
| ZNF385D-AS2 | ZNF717 | ABCA1 | ABCC1 | ABCC4 | ACO1 | ACYP2 | ADAMTS12 | ADAMTS14 | ADCY9 |
| ADGRA3 | ADGRD1 | ADGRV1 | AGAP1 | AK3 | AK9 | AKR1C2 | ALOX12P2 | ANK2 | ANKRD18B |
| ANKS1B | ANO2 | AOPEP | **ARHGEF28** | ASB3 | ATOH1 | ATP2B2 | ATP8A1 | ATP9B | BBS9 |
| **BCL11B** | BORCS5 | BRD1 | C7 | C9orf62 | CACNB2 | CAMK4 | CARMIL1 | CASC20 | CCDC113 |
| CCDC173 | CCNG1 | CCR3 | CD180 | CDC42BPA | CDCA2 | **CDH13** | CDH4 | CHST9 | CNTN5 |
| CNTNAP5 | COA1 | COX18 | CRIM1-DT | CSGALNACT1 | CTNNA3 | CUBN | DAOA-AS1 | DHRS2 | DLEU7-AS1 |
| DLGAP1 | DOCK7 | DOK5 | DPP10 | DSCAM | DSCAML1 | EGF | ENOX1 | EPHA5 | EPHA7 |
| ERCC4 | ESR1 | EXO1 | FAM126A | FAM135B | FAM155A | FBN2 | FCHO1 | FGFR2 | FLJ20021 |
| FNDC3B | FXN | FYN | GABRA5 | GABRB2 | GABRG1 | GAS7 | GLI3 | GLRX3 | GNPDA2 |
| GPR139 | GRM7 | GRM7-AS3 | GRM8 | GTSCR1 | GUSBP1 | HAND2-AS1 | HELT | IL20RB | INPP4B |
| IPO9 | IQGAP2 | **ITPR2** | JAKMIP3 | KALRN | KCNG1 | KCNIP4 | KCNJ6 | KCNQ5 | KCTD16 |
| KIAA1217 | KIAA1549 | KLF4 | KLHL29 | LARS2 | LINC00355 | LINC00430 | LINC00587 | LINC00693 | LINC00882 |
| LINC00885 | LINC01139 | LINC01179 | LINC01192 | LINC01307 | LINC01320 | LINC01419 | LINC01470 | LINC01492 | LINC01493 |
| LINC01537 | LINC01550 | LINC01847 | LINC01933 | LINC01934 | LINC02005 | LINC02070 | LINC02120 | LINC02155 | LINC02211 |
| LINC02233 | LINC02301 | LINC02304 | LINC02315 | LINC02353 | LINC02392 | LINC02405 | LINC02426 | LINC02479 | LINC02494 |
| LINC02508 | LINC02572 | LINC02582 | LINC02740 | LINC02760 | LINS1 | LOC100287387 | LOC100420587 | LOC100505797 | LOC100506207 |
| LOC100507557 | LOC101926960 | LOC101927078 | LOC101927709 | LOC101927815 | LOC101927829 | LOC101927881 | LOC101927967 | LOC101928782 | LOC101928858 |
| LOC101929710 | LOC102546299 | LOC105374704 | LOC105374960 | LOC105374972 | LOC105377448 | LOC105378146 | LOC286178 | LOC643542 | LOC730100 |
| LRRC38 | LRRTM4 | LSM14A | LTO1 | LUZP2 | MAGI3 | MANEA-DT | MAT2B | MBOAT1 | METTL4 |
| MGAT4C | MGC4859 | MIR2054 | MIR302F | MIR3924 | MIR4454 | MIR4539 | MIR4689 | MIR8081 | MRGPRX4 |
| MTHFD1L | MTMR7 | MTPAP | MYO16 | MYO1D | MYO3B | NAALADL2-AS2 | NBEA | NBPF3 | NCALD |
| NDUFC2-KCTD14 | NECTIN3-AS1 | NEDD1 | NPAS3 | NRG1 | NRN1 | NTRK2 | OR4E1 | OTOL1 | PAM |
| PARD3 | PARD3B | PCDH15 | PCDH9 | PCSK5 | PDE8A | PDGFB | PDLIM5 | PLEKHA5 | POLR1D |
| POTEKP | POU6F2 | PRMT8 | PRR20B | PRR5-ARHGAP8 | PRRX1 | PTPRM | PTPRT | QRFPR | RAB9BP1 |
| RAD51B | RASGEF1A | RASSF9 | RBM46 | RBMS3-AS3 | RELN | RWDD4 | S100A10 | SCG2 | SDK1 |
| SDR16C6P | SEMA3C | SETBP1 | SFRP1 | SGCZ | SH3PXD2B | SH3RF1 | SIM1 | SIPA1L1 | SLC16A7 |
| SMOC2 | SMYD3 | SNCAIP | SPOCK3 | STK3 | STK33 | STPG4 | SVIL | TAF4B | TBC1D22A |
| TCTE3 | TDRD15 | TENM3 | TET2 | THORLNC | TMC3-AS1 | TMCO5A | TMEM132C | TMEM200A | TMTC1 |
| TNKS | TRPM3 | TSHR | TSHZ2 | UBE2K | UBE2V2 | UGT8 | UTRN | VENTXP7 | VPS13B |
| VPS53 | VPS8 | WDFY3-AS2 | WWOX | XPNPEP1 | YWHAQ | YY1P2 | ZBTB44 | ZMAT4 | ZMIZ1-AS1 |
| ZNF438 | ZNF890P | ZNRF1 | AAK1 | ABI3BP | ABR | ADGRL3-AS1 | AGBL1 | AKAP13 | ALG8 |
| ALK | AMPH | ANGPT1 | ANK3 | ANKRD34C-AS1 | ANO4 | APTX | ARFGAP3 | ARHGAP24 | ARHGAP8 |
| ARSJ | BCKDHB | BRCA1 | BRINP3 | BTBD16 | C4orf33 | **C9orf72** | CASC16 | CASC6 | CCDC162P |
| CDKN2B-AS1 | CHL1 | CLDN10 | CLPB | CNOT4 | CNTNAP4 | COL23A1 | CPNE4 | CPSF2 | CSMD3 |
| CSNK2A2 | CST7 | DCAF5 | DDX10 | DLC1 | DLEU7 | DMRT1 | DUSP12 | EBF2 | EDIL3 |
| EFNB2 | EPHB1 | EPM2A | ESRRG | EXD2 | EYS | FAM171A1 | FAM181B | FAM227B | FAT3 |
| FGF10 | FH | FMNL2 | FOXN3 | GABRG3 | GALNT13 | GALNT16 | GOLGA2P6 | GRID1 | GRIN2A |
| GRIP1 | HACE1 | HSPA9 | HTR1B | HTR4 | IFNAR2 | IQCB1 | IRAK1BP1 | IRF8 | ISX-AS1 |
| ITGA1 | IWS1 | KCCAT198 | KIRREL3 | KY | LINC00290 | LINC00378 | LINC00461 | LINC00559 | LINC00583 |
| LINC00676 | LINC00910 | LINC00972 | LINC01242 | LINC01262 | LINC01317 | LINC01322 | LINC01350 | LINC01351 | LINC01435 |
| LINC01507 | LINC01520 | LINC01541 | LINC01568 | LINC01592 | LINC01630 | LINC01714 | LINC01718 | LINC01790 | LINC01950 |
| LINC02024 | LINC02064 | LINC02065 | LINC02112 | LINC02143 | LINC02171 | LINC02227 | LINC02346 | LINC02490 | LINC02511 |
| LINC02534 | LINC02605 | LOC100129603 | LOC100288254 | LOC100505501 | LOC100506403 | LOC100506858 | LOC101927237 | LOC101927394 | LOC101927668 |
| LOC101929268 | LOC102724084 | LOC105374060 | LOC283856 | LOC93463 | LPCAT1 | LPP | LRMDA | LRRC9 | LSAMP |
| MACC1 | MDGA2 | MGAT5 | MIR2113 | MIR378C | MIR4275 | MIR8054 | MITF | NBR1 | NME7 |
| NPTX2 | NR2F1-AS1 | NSUN2 | OLFM2 | OPCML | PATJ | PCNX1 | PHIP | PI15 | PLCH1 |
| PLSCR4 | PLXNA2 | POM121L12 | PPIL6 | PPM1H | PRKG1 | PROC | PROX1-AS1 | PRR16 | PTPRD-AS2 |
| PTPRG | RAB20 | RAB3GAP2 | RAN | RBFOX1 | RBPJ | RBPMS-AS1 | RUNX1T1 | SEMA6D | SH2D4A |
| SIN3B | SLC14A2 | SLC35F4 | SLC66A1L | SLC9A9 | SNHG5 | SNORD3G | SORBS2 | SOSTDC1 | SPTSSA |
| ST6GALNAC5 | STAM2 | STPG2 | SURF6 | **SYNE1** | TAMM41 | TAPT1-AS1 | TBX3 | TECRL | TEX41 |
| TMPRSS5 | TMX3 | TPK1 | UBE2E2 | ULK4 | VAV2 | WDR72 | WIPF1 | XKR4 | XYLT1 |
| ZBTB20 | ZIC4 | ZNF121 | ZNF25 | ZNF385D | ZNF426 | AGTPBP1 | APP | AQR | CALN1 |
| CCSER1 | DNAH5 | IQSEC1 | LINC00903 | LOC728755 | MIR4300HG | MSC-AS1 | MYRFL | PABPC4L | PCDH10 |
| PRDM5 | PTPRD | RAB12 | RAX | **SOX5** | ST6GAL1 | TCERG1L | ZNF677 | MAN1A1 | PCSK2 |
| PDE10A | PRKN | | | | | | | | |

**Supplementary Table 4.** 922 genes decisive for ALS classification. 13 genes which have intersection with ALSoD genes are marked as bold.

| | | | | | | | | | |
|---|---|---|---|---|---|---|---|---|---|
| ABAT | ABCG8 | ACCS | ACSL1 | ACVRL1 | ADGRF1 | ADGRG6 | ADGRL4 | **AGT** | AGTR1 |
| AHSA2P | AK2 | ALDH1A3 | AMTN | AMY1C | ANKHD1 | ANKK1 | ANKRD1 | ANKRD16 | ANKS4B |
| ANO2 | AP4B1 | APBB1 | AQP10 | AREL1 | ARG1 | ARIH1 | ARMC2-AS1 | ARNTL2 | ASB18 |
| ASS1 | ATP7B | BCAT1 | BCDIN3D-AS1 | BEND3 | BLACAT1 | BRINP2 | BRMS1L | BTN3A2 | C11orf21 |
| C16orf58 | C16orf97 | C1orf220 | C1QTNF7 | C3orf38 | C5orf30 | C6orf201 | CACNA1E | CACNA2D3-AS1 | CADM2 |
| CADM2-AS2 | CALCR | CALHM1 | CAMKK2 | CAPN10 | CAPN2 | CARNMT1-AS1 | CARS2 | CASP1 | CASP12 |
| CBSL | CBX6 | CC2D2A | CCBE1 | CCDC181 | CCDC18-AS1 | CCDC28A | CCDC86 | CCNB2 | CCNT1 |
| CD207 | CD8B2 | CDC73 | CDH17 | CDKN2A | CDS1 | CELF3 | CFTR-AS1 | CHCHD3 | CHMP1A |
| CHODL | CHST8 | CISTR | CKAP2L | CKLF-CMTM1 | CLYBL-AS1 | CMAHP | CNNM4 | CNP | CNPY3-GNMT |
| CNTN2 | COA7 | COMP | COX10-AS1 | COX7C | CPEB3 | CPT2 | CR1 | CREB3L1 | CREG1 |
| CRISPLD2 | CRSP8P | CRTAC1 | CRTC3 | CRYAB | CRYL1 | CTAGE10P | CTDNEP1 | CTNNAL1 | CUL9 |
| CUTC | CYB5RL | CYP20A1 | CYTIP | CZ1P-ASNS | DAB2 | DBF4 | DCAF1 | DCANP1 | DCDC2 |
| DCHS1 | DHCR24 | DIRC3-AS1 | DKFZp434L192 | DLG1-AS1 | DLGAP1-AS5 | DMRTA2 | DNAAF3 | DNAH8 | DNAJC17 |
| DOCK9-DT | DPF2 | DPF3 | DRAM2 | DUSP5 | E2F7 | EBAG9 | EBLN3P | ECEL1P2 | ECHDC1 |
| EEF1G | EFTUD2 | EIF4ENIF1 | ELFN2 | EMILIN3 | ENTPD6 | EP300 | EPHA10 | EPHA3 | ERP29 |
| ERVV-1 | ETV6 | EVX1 | EYA2 | EZH2 | F11R | F12 | F5 | FAM107B | FAM169A |
| FAM25A | FAM3D-AS1 | FANCM | FBXO15 | FBXO21 | FCF1P2 | FEZF1-AS1 | FKBP3 | FLJ42351 | FNTB |
| FOXB1 | FRAT2 | FTCD | FUBP1 | FYB2 | GAD1 | GAD2 | GALNT18 | GALNT8 | GANC |
| GAS1RR | GATAD2B | GBP5 | GDF7 | GNA14-AS1 | GNAS-AS1 | GNG5 | GNLY | GOLGA8A | GRIK5 |
| GRXCR2 | GTF2B | GTF2F2 | GUSBP2 | GYPC | H3F3A | HCP5B | HDHD5-AS1 | HDLBP | HELQ |
| HIF3A | HOMER1 | HSDL1 | HSH2D | HSPA1B | IFI27L2 | IFNA8 | IFT172 | IFT27 | IGSF11 |
| IGSF9B | IL13 | IMP3 | IMPDH2 | INTS4 | IPO7 | ISCA1 | ITPKA | KANK1 | KAT6A |
| KCNE1B | KCTD11 | KIAA0556 | KIAA2012-AS1 | KIF2A | KIF7 | KLF18 | KLHDC7A | KLHL38 | KLHL42 |
| KLK3 | KMT5A | KRT23 | KRT31 | KRTAP11-1 | KRTAP19-3 | KRTAP19-8 | KRTAP4-5 | KRTAP4-8 | LAMA4 |
| LANCL2 | LBX1-AS1 | LDB1 | LEF1 | LINC00229 | LINC00271 | LINC00272 | LINC00331 | LINC00433 | LINC00536 |
| LINC00562 | LINC00682 | LINC00689 | LINC00705 | LINC00867 | LINC00896 | LINC00954 | LINC01003 | LINC01013 | LINC01122 |
| LINC01215 | LINC01235 | LINC01260 | LINC01354 | LINC01397 | LINC01414 | LINC01424 | LINC01467 | LINC01512 | LINC01515 |
| LINC01531 | LINC01686 | LINC01739 | LINC01744 | LINC01763 | LINC01782 | LINC01865 | LINC01926 | LINC01985 | LINC02022 |
| LINC02069 | LINC02101 | LINC02126 | LINC02162 | LINC02190 | LINC02220 | LINC02228 | LINC02259 | LINC02263 | LINC02324 |
| LINC02333 | LINC02343 | LINC02355 | LINC02362 | LINC02365 | LINC02387 | LINC02388 | LINC02424 | LINC02465 | LINC02471 |
| LINC02475 | LINC02476 | LINC02529 | LINC02574 | LINP1 | LIPA | LOC100130992 | LOC100505635 | LOC100505915 | LOC100507065 |
| LOC100996750 | LOC101926898 | LOC101927394 | LOC101928068 | LOC101928273 | LOC101928708 | LOC101929058 | LOC101929315 | LOC101929427 | LOC101929431 |
| LOC101929529 | LOC101929555 | LOC101929592 | LOC102724452 | LOC105372633 | LOC154449 | LOC283856 | LOC284240 | LOC400499 | LOC400682 |
| LOC441455 | LOC442028 | LOC646626 | LOC729080 | LOC729218 | LOC730101 | LPXN | LRIF1 | LRP1B | LRRC3B |
| LRRC47 | LRRCC1 | LRTM1 | LTK | LUADT1 | LYPD2 | MAB21L4 | MAGI2-AS2 | MAGOH | MAMDC2 |
| MANEA | MAP1LC3A | MAP9 | MAST2 | MCCC2 | MCUR1 | MECR | MEI4 | MEP1A | METTL14 |
| METTL21C | MFSD14A | MGAM | MGP | MICOS10-NBL1 | MIER1 | MIR181C | MIR29B2CHG | MIR3169 | MIR3660 |
| MIR4293 | MIR4464 | MIR4733 | MIR4790 | MIR5580 | MIR8065 | MLLT11 | MMP27 | MMP3 | MPEG1 |
| MPP6 | MPV17L | MSH4 | MSH6 | MTHFD2L | MTSS2 | MTUS1 | MYH7B | MYLK2 | MYORG |
| MYRFL | MYT1L | N6AMT1 | NAA60 | NABP1 | NAT14 | NCAPD3 | NCF4 | NEUROD4 | NFIA-AS2 |
| NGDN | NGLY1 | NIPSNAP3A | NLN | NMI | NOTCH3 | NPFFR1 | NTNG1 | NWD2 | OAS1 |
| OR10H2 | OR10T2 | OR10W1 | OR13C5 | OR1J2 | OR2B11 | OR51A7 | OR52B2 | OR6N2 | OR7E37P |
| OR8B4 | PARAL1 | PAX3 | PAX5 | PBX4 | PCAT5 | PCDHB9 | PCDHGA4 | PCM1 | PDCD4 |
| PEAK1 | PECR | PENK | PFDN2 | PHKG1 | PHOSPHO2-KLHL23 | PI4KB | PIGB | PKD1L2 | PLEKHB1 |
| PLEKHD1 | PLEKHF1 | PLEKHO1 | PMP22 | PODXL | POU3F1 | PPFIA4 | PPP1R13L | PPP1R18 | PPP1R26 |
| PRDM2 | PRDM6 | PRG4 | PRICKLE2-AS3 | PRLH | PRPF40A | PRSS56 | PRSS58 | PSMB3 | PSTK |
| PTGES3L-AARSD1 | PTX3 | **PVR** | QKI | QTRT2 | RAD21-AS1 | RAD51 | RASGRF2-AS1 | RASSF8 | RASSF9 |
| RBPMS | RBSN | RCBTB1 | RCE1 | RDM1 | RDM1P5 | RFLNA | RFT1 | RHOBTB2 | RHOH |
| RHPN1 | RICTOR | RILP | RIMS2 | RING1 | RMND1 | RNF121 | RNF141 | RNF169 | RNF41 |
| RPL13AP5 | S100PBP | SALRNA3 | SATB2 | SBF2-AS1 | SBNO1 | SCGB3A1 | SCN8A | SEC14L4 | SEC62 |
| SELENON | SEPTIN10 | SERPINA11 | SERPINA13P | SERPINA3 | SERPINA4 | SFSWAP | SGO1 | SH3GLB1 | SHH |
| SIAH1 | SIX1 | SLC12A3 | SLC13A4 | SLC16A12 | SLC22A23 | SLC25A30 | SLC2A1-AS1 | SLC30A8 | SLC38A7 |
| SLC41A2 | SLC44A2 | SLC46A1 | SLC46A3 | SLC48A1 | SLC66A3 | SLC9B1 | SLCO2A1 | SMC5 | SMIM35 |
| SMIM7 | SMR3A | SMUG1 | SNAPC1 | SNAR-A12 | SNORA33 | SNORD3J | SNRK-AS1 | SNX3 | SOBP |
| SP110 | SPACA6P-AS | SPATA17 | SPATA3-AS1 | SPATA48 | SPDEF | SPEG | SPIB | SPINT1 | SPTBN5 |
| SPTLC3 | SRGAP2 | SRRM1 | SRSF6 | SSX2IP | ST8SIA1 | STAG3L5P-PVRIG2P-PILRB | STAT3 | STIL | STK4 |
| STMP1 | STOX2 | STXBP5 | SUSD6 | SYCP3 | SZT2 | TCTE3 | TEMN3-AS1 | TENT2 | TEX12 |
| TEX52 | TEX53 | TFCP2 | THOC5 | TIMM44 | TLE7 | TLX1NB | TMBIM6 | TMED8 | TMEM161B-AS1 |
| TMEM212-AS1 | TMEM233 | TMEM246 | TMEM254-AS1 | TMPRSS11F | TMPRSS7 | TPT1-AS1 | TRAF6 | TRIM2 | TRIM58 |
| TRIM71 | UBLCP1 | UHRF2 | UNC119B | UNC80 | USE1 | USH1C | USHBP1 | USP18 | USP39 |
| VAV1 | VCAM1 | VEZT | VSNL1 | WAPL | WDR92 | XIRP1 | XPOT | YRDC | YTHDF3-AS1 |
| ZBTB16 | ZBTB40 | ZC3HC1 | ZDHHC18 | ZDHHC19 | ZEB1-AS1 | ZFAND5 | ZFP2 | ZFPM2-AS1 | ZIM2-AS1 |
| ZKSCAN5 | ZNF136 | ZNF143 | ZNF23 | ZNF284 | ZNF318 | ZNF32-AS3 | ZNF33B | ZNF341 | ZNF385A |
| ZNF396 | ZNF426-DT | ZNF445 | ZNF480 | ZNF492 | ZNF570 | ZNF597 | ZNF606 | ZNF704 | ZNF705E |
| ZNF804A | ZNF829 | ZSCAN5A | ZSWIM9 | | | | | | |

**Supplementary Table 5.** Potential non-additive genes associated with ALS. Two genes which have intersection with ALSoD genes are marked as bold.

| Batch identifier | #Case(ALS) | #Control(Healthy) | Total |
|---|---|---|---|
| C1 | 225 | 380 | 605 |
| C3 | 130 | 49 | 179 |
| C5 | 0 | 5155 | 5155 |
| C44 | 2837 | 1629 | 4466 |
| Total | 3192 | 7213 | 10405 |

**Supplementary Table 6.** Batch structure of our data set. Values in the table indicate the number of samples.

| ***Precision*** | | | | | | |
|---|---|---|---|---|---|---|
| DR | Gene-PCA | Gene-PCA | Gene-PCA | Gene-PCA | Gene-PCA | - |
| Classifiers | SVM | MLP | CNN | DiseaseCapsule | LR | LR* |
| 5% | *nan* | $84.7 \pm 2.3$ | $60.9 \pm 7.3$ | $74.2 \pm 4.7$ | $64.9 \pm 0.5$ | $94.0 \pm 1.4$ |
| 10% | $100.0 \pm 0.0$ | $90.2 \pm 1.4$ | $68.2 \pm 3.8$ | $79.8 \pm 2.8$ | $66.9 \pm 0.3$ | $92.1 \pm 1.5$ |
| 20% | $96.8 \pm 0.1$ | $91.3 \pm 0.9$ | $80.5 \pm 3.7$ | $81.5 \pm 1.5$ | $67.0 \pm 0.3$ | $90.7 \pm 1.6$ |
| 40% | $94.7 \pm 0.7$ | $91.4 \pm 0.9$ | $82.9 \pm 2.0$ | $82.6 \pm 1.2$ | $68.0 \pm 0.3$ | $91.3 \pm 1.0$ |
| 60% | $94.2 \pm 0.2$ | $91.9 \pm 1.0$ | $84.7 \pm 1.7$ | $83.0 \pm 1.0$ | $69.0 \pm 0.3$ | $91.0 \pm 0.9$ |
| 80% | $93.5 \pm 0.2$ | $91.6 \pm 0.6$ | $85.6 \pm 1.6$ | $83.4 \pm 0.6$ | $69.7 \pm 0.3$ | $91.0 \pm 0.7$ |
| 100% | $94.8 \pm 0.4$ | $92.8 \pm 0.9$ | $86.1 \pm 1.2$ | $84.6 \pm 0.5$ | $71.6 \pm 0.3$ | $91.6 \pm 0.3$ |
| ***Recall*** | | | | | | |
| DR | Gene-PCA | Gene-PCA | Gene-PCA | Gene-PCA | Gene-PCA | - |
| Classifiers | LR | DiseaseCapsule | MLP | CNN | SVM | LR* |
| 5% | $83.3 \pm 0.4$ | $74.8 \pm 8.1$ | $28.1 \pm 2.6$ | $32.2 \pm 24.6$ | $0.0 \pm 0.0$ | $21.5 \pm 2.4$ |
| 10% | $87.7 \pm 0.6$ | $77.9 \pm 3.8$ | $35.8 \pm 1.5$ | $24.3 \pm 9.8$ | $6.3 \pm 0.5$ | $37.3 \pm 1.4$ |
| 20% | $92.0 \pm 0.4$ | $80.9 \pm 2.9$ | $47.4 \pm 1.1$ | $31.5 \pm 7.0$ | $23.6 \pm 0.5$ | $48.8 \pm 1.1$ |
| 40% | $92.9 \pm 0.2$ | $85.0 \pm 1.8$ | $58.6 \pm 1.3$ | $42.5 \pm 1.8$ | $36.6 \pm 0.4$ | $59.4 \pm 1.6$ |
| 60% | $94.8 \pm 0.3$ | $86.2 \pm 1.3$ | $65.7 \pm 1.3$ | $49.3 \pm 3.0$ | $44.9 \pm 0.3$ | $65.1 \pm 1.1$ |
| 80% | $94.5 \pm 0.2$ | $87.6 \pm 0.8$ | $69.3 \pm 1.2$ | $53.6 \pm 1.7$ | $49.8 \pm 0.5$ | $68.2 \pm 0.8$ |
| 100% | $94.5 \pm 0.2$ | $88.7 \pm 0.4$ | $72.7 \pm 1.2$ | $56.5 \pm 1.3$ | $55.2 \pm 0.4$ | $70.3 \pm 0.6$ |
| ***F1 score*** | | | | | | |
| DR | Gene-PCA | Gene-PCA | Gene-PCA | Gene-PCA | Gene-PCA | - |
| Classifiers | DiseaseCapsule | MLP | LR | SVM | CNN | LR* |
| 5% | $74.1 \pm 2.3$ | $42.1 \pm 2.9$ | $72.9 \pm 0.4$ | *nan* | $35.9 \pm 20.2$ | $35.0 \pm 3.1$ |
| 10% | $78.7 \pm 1.0$ | $51.3 \pm 1.7$ | $75.9 \pm 0.4$ | $11.8 \pm 0.9$ | $34.8 \pm 8.8$ | $53.0 \pm 1.4$ |
| 20% | $81.2 \pm 0.9$ | $62.3 \pm 1.1$ | $77.5 \pm 0.3$ | $37.9 \pm 0.7$ | $44.7 \pm 6.8$ | $63.5 \pm 1.1$ |
| 40% | $83.7 \pm 0.6$ | $71.4 \pm 1.0$ | $78.5 \pm 0.3$ | $52.7 \pm 0.5$ | $56.2 \pm 1.6$ | $72.0 \pm 1.3$ |
| 60% | $84.6 \pm 0.3$ | $76.6 \pm 1.0$ | $79.8 \pm 0.3$ | $60.8 \pm 0.3$ | $62.3 \pm 2.5$ | $75.9 \pm 0.8$ |
| 80% | $85.5 \pm 0.4$ | $78.9 \pm 0.9$ | $80.2 \pm 0.2$ | $65.0 \pm 0.4$ | $65.9 \pm 1.5$ | $78.0 \pm 0.7$ |
| 100% | $86.6 \pm 0.3$ | $81.6 \pm 1.0$ | $81.5 \pm 0.2$ | $69.7 \pm 0.3$ | $68.2 \pm 1.0$ | $79.5 \pm 0.4$ |

**Supplementary Table 7.** Test precision, recall and F1 score of various models trained using different percentage of subsampling training samples, namely 100%, 80%, 60%, 40%, 20%, 10%, 5%. The first column denotes the subsampling percentages. Values are represented as mean ± sd (%), which are calculated for the same test dataset by repeating the subsampling and training-testing processes for ten times with the identical network architecture and hyperparameters. Columns are ordered by their corresponding metric values. *nan*: this metric cannot be calculated, DR: dimensionality reduction, LR: Logistic Regression. *This model is PRS-based that the SNPs were selected by GWAS with the threshold ($P < 5 \times 10^{-2}$). *Note*: The best value is marked in bold.
